# The sympathetic nervous system initiates liver regeneration

**DOI:** 10.64898/2026.07.16.739048

**Authors:** Hiroshi Nagao, Satoshi Kofuji, Keiko Danzaki, Akira Suzuki, Hiroshi Nishina

## Abstract

Within hours of injury to the liver, signals from damaged hepatocytes activate Kupffer cells, inducing the production of inflammatory cytokines and proliferation of quiescent hepatocytes to regenerate the liver^1,2^. In humans, liver regenerative capacity declines with age^3,4^. However, the nature of the initial signals of liver regeneration and how they are affected during aging remain unclear. In this study, we leveraged “neuro-aging mice”, in which neuronal function declines with age^5^, and discovered an essential role for intrahepatic sympathetic nerves in the early phase of liver regeneration. Upon liver injury in wild-type mice, there was a rapid, transient production of noradrenaline (NA) in the liver. This was abolished in neuro-aging mice who instead exhibited decreased metabolic capacity and fatal liver regeneration failure. NA coordinated the activity of multiple cell types by inducing expression of acute-response genes in hepatocytes and stimulating production of inflammatory cytokines such as IL-6 in Kupffer cells. These results identify a crucial role for the sympathetic nervous system and NA in liver regeneration. Further, they suggest that aging-related loss of this sympathetic nerve-NA regulatory axis reduces liver regenerative capacity.

## Introduction

The liver acts as the primary hub for detoxification and metabolism. In 1931, American surgeons Higgins and Anderson experimentally demonstrated that the liver is an organ capable of regeneration^6^. Since then, for nearly a century, liver regeneration has attracted the attention of numerous biologists and clinicians. Liver regeneration is an orchestrated homeostatic mechanism involving the precise coordination of multiple cell types, including hepatocytes, resident macrophages (Kupffer cells), and hepatic stellate cells. This process is traditionally divided into three distinct phases: the initiation phase (0–24 h), where quiescent (G_0_) hepatocytes transition into the G_1_ phase; the proliferation phase (24–72 h), characterized by active cell division; and the termination phase (beyond 72 h), where regeneration ceases once the original mass is restored^2^. During the initiation phase, Kupffer cells function as central mediators, secreting TNF-α and IL-6 to prime hepatocytes for reentry into the cell cycle. The proliferation phase is driven by HGF and EGF, derived in part from the extracellular matrix, which trigger DNA synthesis and expansion of the hepatocyte population^1^. Finally, regeneration is concluded in the termination phase through TGF-β signaling, which inhibits further proliferation^7^. Numerous studies have established Kupffer cells as central mediators promoting hepatocyte proliferation^8^. However, fundamental questions remain: How is the initial injury sensed within hours? What molecular mechanism coordinates the diverse cell types in the liver to launch the regenerative program? And what causes the decline in liver regenerative capacity with aging?

Several studies have highlighted a role for the parasympathetic nervous system, specifically the vagus nerve, which induces acetylcholine (ACh), in activating Kupffer cells and promoting liver regeneration^9–11^. For example, hepatic vagotomy to eliminate ACh surge or treatment with an ACh antagonist impaired liver regeneration after partial hepatectomy (PHx)^9^. And the addition of ACh to isolated macrophages induced IL-6 production^9^. Recently, it was reported that ACh derived from intrahepatic B cells activates Kupffer cells to induce IL-6 expression^12^. However, it is the sympathetic nervous system that is most densely distributed throughout the liver^13–15^. The sympathetic nervous system induces NA and is considered a responder to physiological changes such as metabolic regulation, inflammation, and vascular tone^16–18^. Early studies showed that circulating NA levels surge following PHx and can promote hepatocyte DNA synthesis^19,20^. However, whether and how the sympathetic nervous system triggers liver regeneration upon injury is currently unclear.

The age-related decline in tissue and organ function is increasingly attributed not only to cell-intrinsic senescence but also to the deterioration of systemic regulatory networks, such as the circulatory and nervous systems^21–23^. For instance, sarcopenia—the geriatric loss of muscle mass and strength—is driven by the progressive dysfunction of the neuromuscular junction^24–27^. Similarly, the liver exhibits a profound decline in metabolic and regenerative capacity with advancing age^28–30^. However, whether dysfunction of the autonomic nervous system drives this impaired regenerative response in the aging liver has yet to be determined. Previously, we established a mature neuron-specific *Mkk7*-deficient mouse model (*Mkk7*^flox/flox^ *Synapsin I-Cre*)^5,31^. In these mice, the phosphorylation of JNK—a key regulator of axonal transport—is reduced by half, leading to accelerated age-related axonal transport defects and a sarcopenia-like phenotype. In the present study, we leveraged this mouse model; hereafter referred to as *Mkk7* cKO or neuro-aging mice, to identify the mechanisms by which neural decline impairs liver regeneration.

## Results

### *Mkk7* cKO mice exhibit age-dependent impairment of liver regeneration

We previously demonstrated that *Mkk7* deletion in our model is specific to mature neurons^31^. To confirm that *Mkk7* remains intact in non-neuronal tissues, we analyzed genomic DNA and protein expression in the liver. Consistent with our previous findings, *Mkk7* was not deleted in the liver of *Mkk7* cKO mice (Extended Data Fig. 1a–d). At 10 months of age, *Mkk7* cKO mice displayed reduced motor coordination due to age-dependent motor neuron dysfunction^5^ (Extended Data Fig. 1e). To investigate the impact of neural aging on liver resilience, we subjected 6-month-old and 10-month-old *Mkk7* cKO mice to PHx or drug-induced liver injury (3,5-diethoxycarbonyl-1,4-dihydrocollidine (DDC), thioacetamide (TAA), or carbon tetrachloride (CCl_4_)). While no significant difference in survival was observed between *Mkk7*^flox/flox^ and *Mkk7* cKO mice at 6 months of age, 10-month-old *Mkk7* cKO mice exhibited decreased survival (Fig. 1a–c). To identify the cause of this age-dependent lethality, we analyzed liver tissues 21 days after CCl_4_ administration, a time point at which 50% mortality was reached (Extended Data Fig. 2a). Serum levels of the liver injury markers ALT and AST were significantly elevated in 10-month-old *Mkk7* cKO mice compared to those in *Mkk7*^flox/flox^ mice and 3- or 6-month-old *Mkk7* cKO mice (Extended Data Fig. 2b). Histological analysis revealed that 10-month-old *Mkk7* cKO mice suffered from exacerbated necrotic areas in the pericentral regions compared to *Mkk7*^flox/flox^ mice. This was accompanied by a nearly complete loss of hepatocyte proliferation (Ki67-positive cells) and a significant reduction in macrophage infiltration (F4/80-positive cells) at the injury sites (Fig. 1d and e).

**Figure 1.**
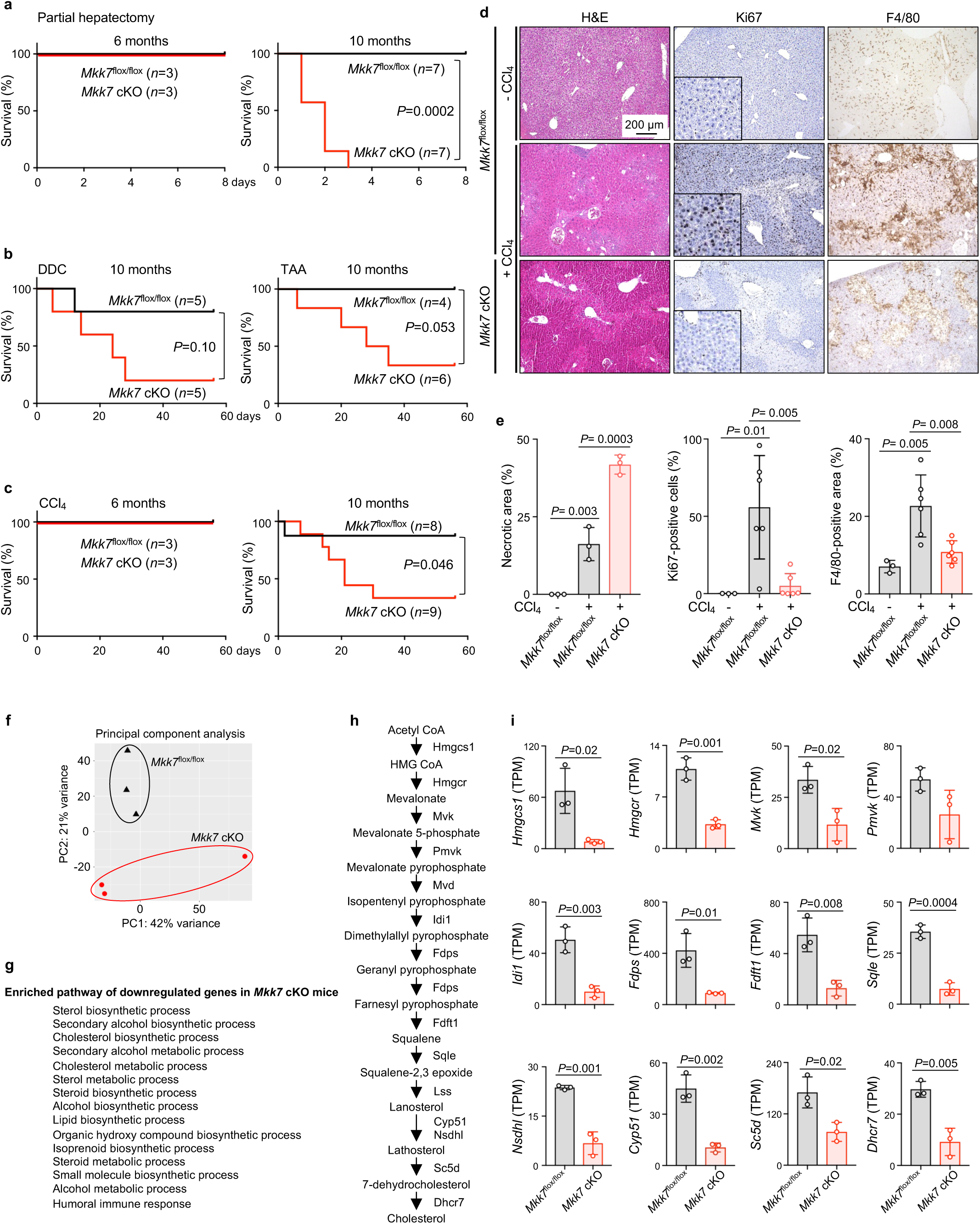
Responses of *Mkk7* cKO mice to liver injury. (**a**) Survival curves of *Mkk7*^flox/flox^ mice and *Mkk7* cKO mice at the indicated ages after PHx. Log-rank test was performed. (**b**) Survival curves of *Mkk7*^flox/flox^ mice and *Mkk7* cKO mice at 10 months of age after DDC and TAA treatment. Log-rank test was performed. (**c**) Survival curves of *Mkk7*^flox/flox^ mice and *Mkk7* cKO mice at the indicated ages after CCl_4_ treatment. Log-rank test was performed. (**d**) H&E staining, anti-Ki67 antibody staining, and anti-F4/80 antibody staining of the liver after chronic CCl_4_ administration. (**e**) Quantification of (d). The data are mean±s.d. *n*=3 mice for each group for H&E staining, *n*=3 for *Mkk7*^flox/flox^ mice (no treatment) and *n*=6 for *Mkk7*^flox/flox^ mice (CCl_4_ treatment) and *Mkk7* cKO mice for anti-Ki67 antibody and anti-F4/80 antibody staining. One-way ANOVA with Tukey multiple comparisons test was performed. (**f**) The principal component analysis of RNA-seq analysis of the livers of *Mkk7*^flox/flox^ mice and *Mkk7* cKO mice at 10 months of age after chronic CCl_4_ administration. *n*=3 mice for each group. (**g**) Enrichment analysis of differentially downregulated genes (Cluster F in Extended Data Fig.2c) in *Mkk7* cKO mice. (**h**) Overview of the sterol biosynthetic pathway. (**i**) Expression of genes regulating the sterol biosynthetic pathway from RNA-seq analysis. The data are mean±s.d. *n*=3 mice for each group. Unpaired two-tailed Student *t* test was performed.

To further elucidate the molecular basis for the compromised hepatocyte survival and proliferation, we performed RNA-sequencing (RNA-seq) on liver samples. Principal component analysis (PCA) showed a distinct separation between 10-month-old *Mkk7*^flox/flox^ and *Mkk7* cKO livers (Fig. 1f). Heatmap and Gene Ontology (GO) enrichment analyses revealed that genes associated with the sterol biosynthesis pathway were downregulated in the *Mkk7* cKO livers (Fig. 1g–i, Extended Data Fig. 2c, Supplementary Table 1). Collectively, these results suggest that 10-month-old *Mkk7* cKO mice undergo a metabolic decline that impairs hepatocyte survival and proliferation, ultimately leading to impaired liver regeneration and death.

### In *Mkk7* cKO mice, aging leads to the loss of neurotransmitter production induced by liver injury

To identify the mechanisms underlying this age-dependent regenerative failure in *Mkk7* cKO (neuro-aging) mice, we first characterized intrahepatic sympathetic innervation. Utilizing tissue clearing technology combined with sympathetic-specific immunostaining, we observed that while thick sympathetic nerve trunks remained detectable in the central region of 13-month-old *Mkk7* cKO livers, peripheral sympathetic fibers were significantly denervated compared to those in *Mkk7*^flox/flox^ mice (Fig. 2a–c, Extended Data Fig. 3a). We next assessed whether this structural decline translated into functional impairment by measuring intrahepatic noradrenaline (NA) levels. In 6-month-old *Mkk7*^flox/flox^ and *Mkk7* cKO mice, CCl_4_ administration triggered a fairly rapid and transient surge in NA, beginning at 3 h and peaking at 6 h post-CCl_4_ (Fig. 2d). In contrast, intrahepatic acetylcholine (ACh) levels, representing parasympathetic activity, peaked later, at 12 h (Fig. 2e). Remarkably, the injury-induced surges of both NA and ACh were completely abolished in the regeneration impaired 10-month-old *Mkk7* cKO mice (Fig. 2f, g). A similar decline in NA and ACh induction at 6 h was observed in 24-month-old naturally aged wild-type mice (Fig. 2f right and g right). To investigate whether there are sex differences in the injury-induced surges of both NA and ACh, female mice were used. CCl_4_ administration induced intrahepatic surges of both NA and ACh in 8-week-old female wild-type mice in the same time course as male mice (Extended Data Fig. 3b). In contrast, the injury-induced surges of both NA and ACh were completely abolished in 13-month-old *Mkk7* cKO female mice (Extended Data Fig. 3c). Thus, there is no sex differences in the injury-induced surges of either NA or ACh (Extended Data Fig. 3d). To determine whether the damage-induced NA was produced in the liver or derived from the systemic circulation, we measured NA levels in the adrenal glands, the primary site of systemic NA production. In 10-month-old *Mkk7*^flox/flox^ mice, adrenal NA levels increased persistently from 3 to 24 h post-CCl_4_, reaching concentrations approximately five times higher than those measured in the liver—a transient NA peak in the liver versus a sustained increase in the adrenal glands (Fig. 2f and h). However, this adrenal NA induction was absent in 10-month-old *Mkk7* cKO mice (Fig. 2h). To test if the sympathetic NA response depends on parasympathetic signaling, we performed hepatic vagotomy (HV) (Fig. 2i). While HV eliminated the ACh surge, the NA surge remained intact (Fig. 2j and k). Similar neurotransmitter dynamics were observed following PHx, with NA and ACh levels peaking 3 h post-surgery in 9-week-old wild-type mice (Fig. 2l and m). Collectively, these data indicate that liver injury triggers a rapid, transient, and hepatic induction of NA and ACh via the central nervous system during the initiation phase (within hours) of liver regeneration (Extended Data Fig. 4). The NA surge is neither a result of adrenal efflux nor dependent on vagal activity. Notably, the hepatic NA and ACh surges were lost upon both artificially (*Mkk7* cKO mice) and naturally (24-month-old wild-type mice) induced aging.

**Figure 2.**
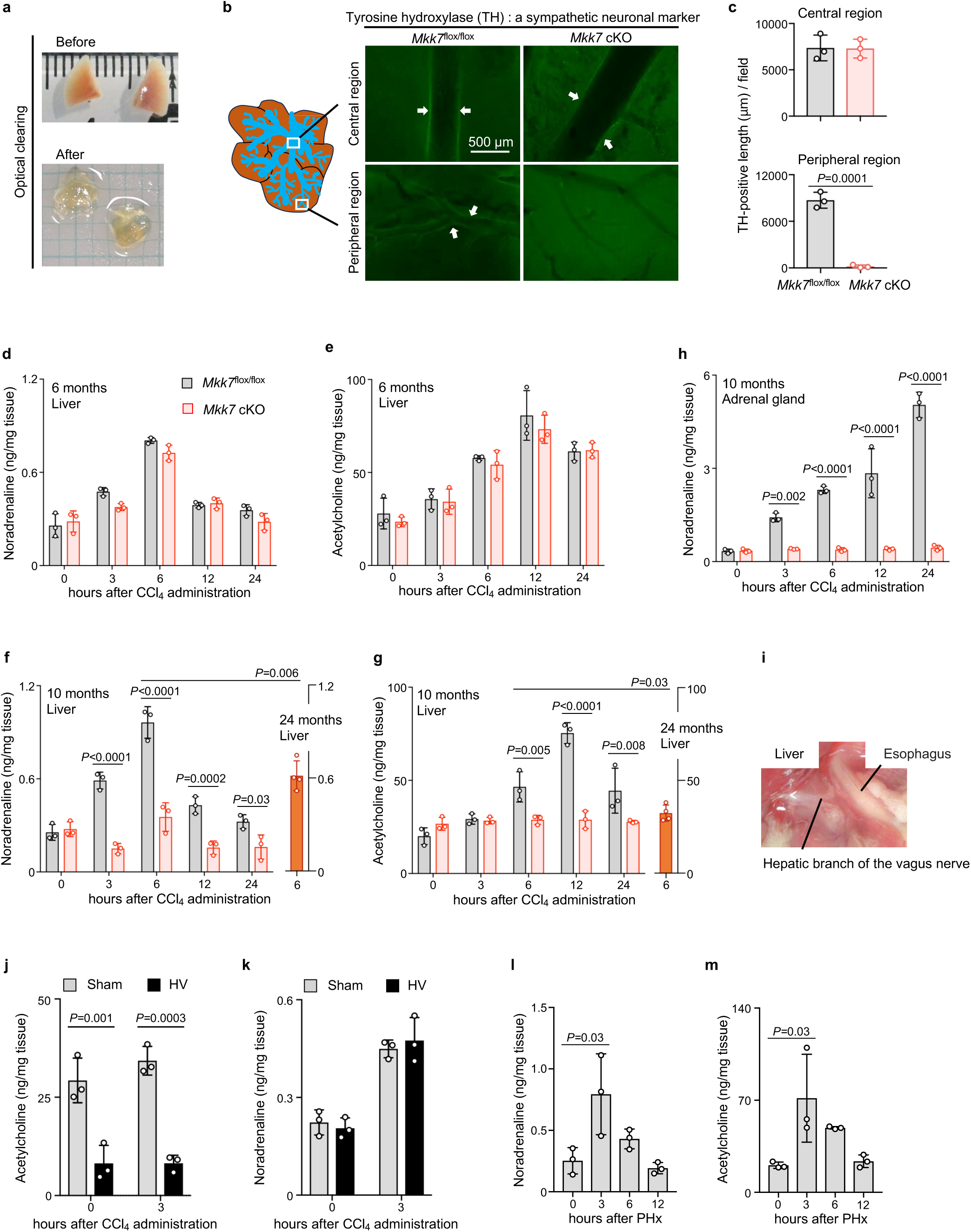
Autonomic nervous responses of *Mkk7* cKO mice to CCl_4_ injury. (**a**) Examples of liver tissue clearing. (**b**) Staining of tyrosine hydroxylase in the left lobe of the liver from mice at 13 months of age. White arrows indicate neural fibers. (**c**) Quantification of neural lengths. The data are mean±s.d. *n*=3 mice for each group. Unpaired two-tailed Student *t* test was performed. (**d**) Measurement of NA levels in the livers of *Mkk7*^flox/flox^ mice and *Mkk7* cKO mice at 6 months of age after CCl_4_ administration. The data are mean± s.d. *n*=3 mice for each group. Two-way ANOVA with Sidak’s multiple comparisons test was performed. (**e**) Measurement of ACh levels in the livers of *Mkk7*^flox/flox^ mice and *Mkk7* cKO mice at 6 months of age after CCl_4_ administration. The data are mean±s.d. *n*=3 mice for each group. Two-way ANOVA with Sidak’s multiple comparisons test was performed. (**f**) Measurement of NA levels in the livers of *Mkk7*^flox/flox^ mice and *Mkk7* cKO mice at 10 months of age and wild-type mice at 24 months of age after CCl_4_ administration. The data are mean±s.d. *n*=3 for *Mkk7*^flox/flox^ and *Mkk7* cKO mice, and *n*=4 for wild-type mice. Two-way ANOVA with Sidak’s multiple comparisons test was performed. For the comparison of the 6 h timepoint between 10 months of age and 24 months of age, unpaired two-tailed Student *t* test was performed. (**g**) Measurement of ACh levels in the livers of *Mkk7*^flox/flox^ mice and *Mkk7* cKO mice at 10 months of age and wild-type mice at 24 months of age after CCl_4_ administration. The data are mean±s.d. *n*=3 for *Mkk7*^flox/flox^ and *Mkk7* cKO mice, and *n*=4 for wild-type mice. Two-way ANOVA with Sidak’s multiple comparisons test was performed. For the comparison of the 6 h timepoint between 10 months of age and 24 months of age, unpaired two-tailed Student *t* test was performed. (**h**) Measurement of NA levels in the adrenal glands of *Mkk7*^flox/flox^ mice and *Mkk7* cKO mice at 10 months of age after CCl_4_ administration. The data are mean±s.d. *n*=3 mice for each group. Two-way ANOVA with Sidak’s multiple comparisons test was performed. (**i**) A photograph of the hepatic branch of the vagus nerve. Measurement of (**j**) ACh and (**k**) NA levels in the wild-type liver at 15 weeks of age after CCl_4_ treatment. The data are mean±s.d. *n*=3 mice for each group. Two-way ANOVA with Tukey’s multiple comparisons test was performed. Measurement of levels of (**l**) NA and (**m**) ACh in the livers of wild-type mice at 9 weeks of age after PHx. The data are mean±s.d. *n*=3 mice for each group. One-way ANOVA with Tukey’s multiple comparisons test was performed.

### Noradrenaline is necessary and sufficient for hepatocyte protection

We next examined the roles of NA and ACh in liver regeneration during the initiation phase—the window between injury and the onset of regeneration. This phase was classified as 12 h post-CCl_4_, when we observed hepatocyte damage and the formation of macrophage clusters, whereas hepatocyte proliferation (Ki67 staining) was not yet evident (Extended Data Fig. 5). To differentiate the roles of NA and ACh in this initiation phase, we treated wild-type mice with their specific neuronal receptor antagonists along with CCl_4_: α_1_- and β-adrenergic receptor antagonists (prazosin and propranolol, respectively) or a muscarinic ACh receptor antagonist (atropine). CCl_4_ normally promoted macrophage cluster formation after 12 h. Adrenergic blockade significantly inhibited this clustering (Fig. 3a top and b) and dramatically increased the number of Cleaved Caspase-3-positive apoptotic hepatocytes (Fig. 3a bottom and c). In contrast, atropine treatment had no significant effect on either macrophage clustering or hepatocyte apoptosis at this stage (Fig. 3d-f). In wild-type mice, liver injury naturally triggers surges of both NA and ACh (Fig. 2d-g), making it difficult to determine whether NA alone is sufficient to drive the protective phenotype. To isolate the specific contribution of NA, we utilized 10-month-old *Mkk7* cKO mice, in which neither the NA nor the ACh surge occurs (Fig. 2f and g). At 12 h post-CCl_4_, these neuro-aging mice exhibited impaired macrophage cluster formation and a profound increase in Cleaved Caspase-3-positive apoptotic hepatocytes (Fig. 3g-i). Interestingly, addition of NA together with CCl_4_ significantly suppressed Cleaved Caspase-3 levels, although it failed to rescue macrophage cluster formation (Fig. 3g-i). These results suggest that during the initiation phase of regeneration, NA—rather than ACh—serves as the primary regulator of the hepatocyte protective phenotype.

**Figure 3.**
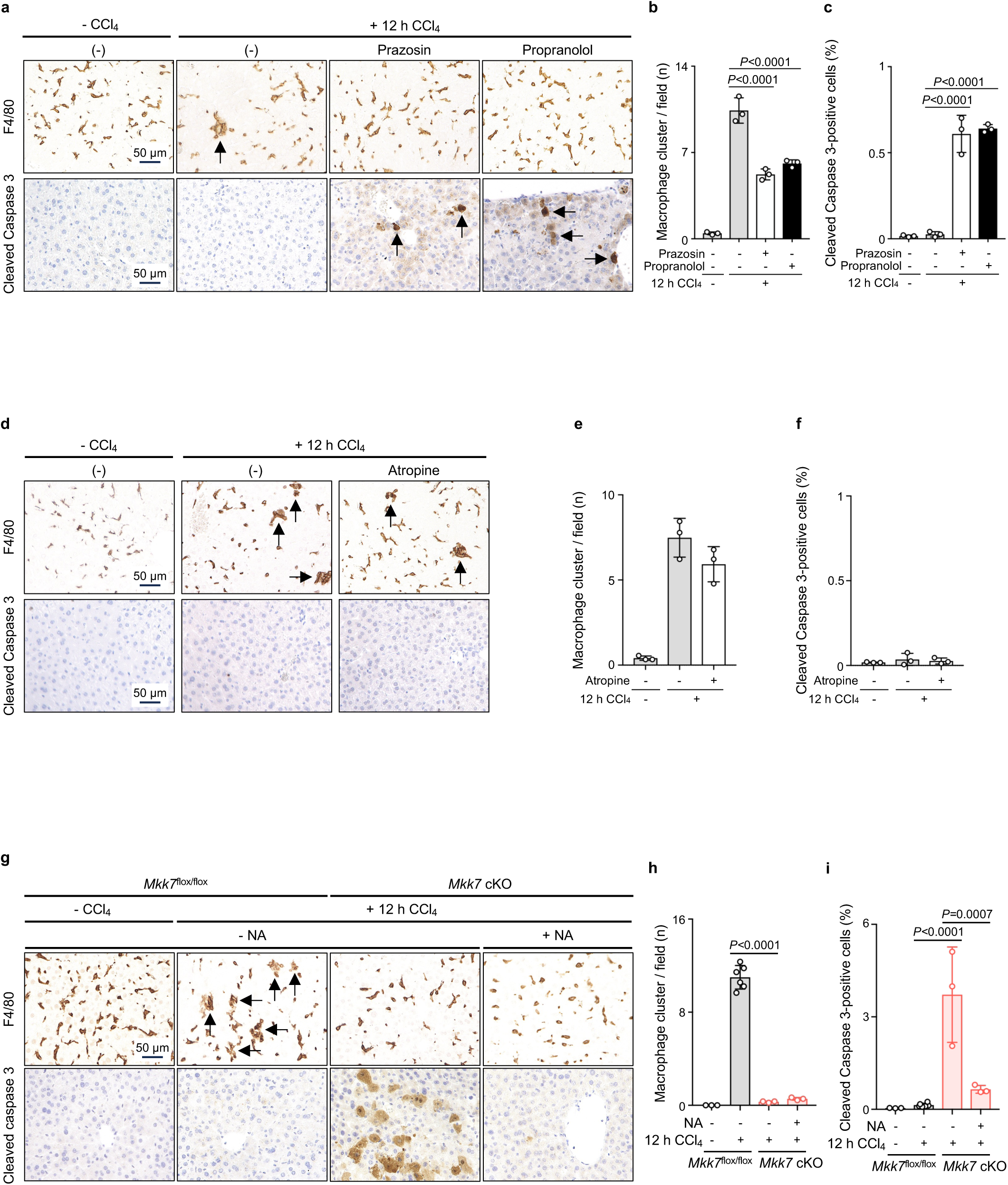
Sympathetic regulation of hepatic cells after CCl_4_ injury. (**a**) Immunostaining of liver sections from wild-type mice at 8 weeks of age treated with CCl_4_ and prazosin or propranolol for 12 h with anti-F4/80 and Cleaved Caspase 3 antibodies. (**b**) Quantification of the number of macrophage clusters from (a). (**c**) Quantification of the ratio of Cleaved Caspase 3-positive cells from (a). The data are mean±s.d. *n*=3 mice for each group. One-way ANOVA with Tukey’s multiple comparisons test was performed. (**d**) Immunostaining of liver sections from wild-type mice at 8 weeks of age treated with CCl_4_ and atropine for 12 h with anti-F4/80 and Cleaved Caspase 3 antibodies. (**e**) Quantification of the number of macrophage clusters from (d). (**f**) Quantification of the ratio of Cleaved Caspase 3-positive cells from (d). The data are mean±s.d. *n*=3 mice for each group. One-way ANOVA with Tukey’s multiple comparisons test was performed. (**g**) Immunostaining of liver sections from *Mkk7*^flox/flox^ mice and *Mkk7* cKO mice at 10 or 11 months of age treated with CCl_4_ and NA for 12 h with anti-F4/80 and Cleaved caspase 3 antibodies. (**h**) Quantification of the number of macrophage clusters from (g). (**i**) Quantification of the ratio of Cleaved Caspase 3-positive cells from (g). The data are mean±s.d. *n*=6 mice for CCl_4_-treated *Mkk7*^flox/flox^ mouse group and *n*=3 mice for the other groups. One-way ANOVA with Tukey’s multiple comparisons test was performed.

### Noradrenaline directly regulates gene expression in hepatocytes and macrophages

Having identified a role for NA in hepatic regeneration, we next sought to identify the molecular mechanisms involved. First, we investigated its potential effects on hepatocytes by performing RNA-seq on liver samples from wild-type mice following systemic administration of NA or an ACh receptor agonist (carbachol), under conditions without CCl_4_ administration. PCA revealed that NA treatment induced a distinct transcriptional profile in hepatocytes within 3 h, which was diminished by 6 and 12 h (Fig. 4a, Extended Data Fig. 6a). Compared to carbachol, NA specifically upregulated genes associated with inflammatory, defense, and acute-phase responses (Fig. 4b, Extended Data Fig. 6b–d, Supplementary Table 2 and 3). Key upregulated genes included *Tlr2* (an innate immune response protein), *Relb* (a noncanonical NF-_K_B protein), *Saa1/2* (acute-phase response proteins), and *S100a8* (a calcium-binding protein). SAA1 binds to TLR2 and activates NF-_K_B signaling^32^. Quantitative PCR (qPCR) confirmed that *Tlr2* and *Relb* induction occurred as early as 2 h post-NA (Extended Data Fig. 7a, b, c), while carbachol had no effect (Extended Data Fig. 7d, e). The induction of these genes was maintained even after Kupffer cell depletion using clodronate, confirming their hepatocyte-specific expression (Extended Data Fig. 7a, b, c, n). We next investigated whether the expression of these genes is upregulated by NA production after liver injury (CCl_4_). We validated that *Tlr2*, *Relb*, *Saa1/2*, and *S100a8* were induced by CCl_4_ in an NA-dependent manner, as their expression was significantly suppressed by the co-administration of prazosin and propranolol (Fig. 4c, d, Extended Data Fig. 8a, b, c, d). Notably, this CCl_4_-NA-dependent gene induction was almost entirely abolished in 10-month-old *Mkk7* cKO mice (Fig. 4e, Extended Data Fig. 8e, f). Together, these results suggest that the sympathetic nervous system (NA) provides a protective effect in the liver by activating a transcriptional program in hepatocytes that mediates inflammatory, defense and acute-phase responses, and that this protective effect is lost during aging.

**Figure 4.**
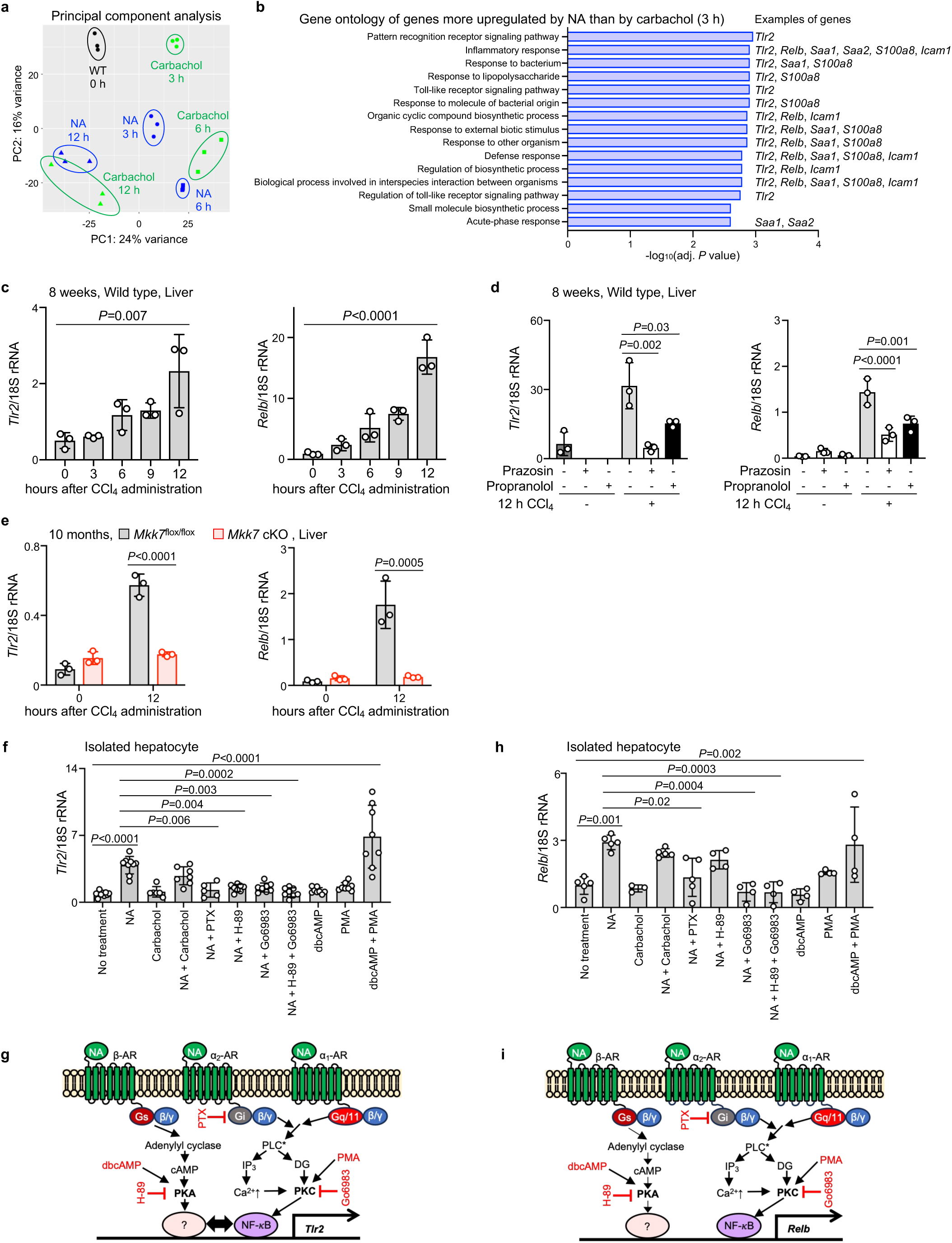
Effects of noradrenaline and carbachol on gene expression in the liver and isolated hepatocytes. (**a**) Principal component analysis of livers from wild-type (WT) mice at 9 or 10 weeks of age treated with NA or carbachol for the indicated times. (**b**) Ontology analysis of genes upregulated by NA compared to carbachol at 3 h after CCl_4_ treatment. Examples of genes that were further investigated in this study are shown on the right. (**c**) Expression of the *Tlr2* gene and the *Relb* gene in the livers from wild-type mice at 8 weeks of age until 12 h after CCl_4_ treatment. The data are mean±s.d. *n*=3 mice for each group. One-way ANOVA with Tukey’s multiple comparisons test was performed. (**d**) Expression of the *Tlr2* gene and the *Relb* gene in the livers from wild-type mice at 8 weeks of age pre-treated with prazosin or propranolol after 12 h treatment of CCl_4_. The data are mean±s.d. *n*=3 mice for each group. One-way ANOVA with Tukey’s multiple comparisons test was performed. (**e**) Expression of the *Tlr2* gene and the *Relb* gene in the livers from *Mkk7*^flox/flox^ mice and *Mkk7* cKO mice at 10 months of age after 12 h treatment of CCl_4_. The data are mean±s.d. *n*=3 mice for each group. Two-way ANOVA with Sidak’s multiple comparisons test was performed. (**f**) qPCR analysis of effects of NA, carbachol, and inhibitors on expression of the *Tlr2* gene in isolated hepatocytes. The data are mean±s.d. *n*=5-9 biologically independent hepatocytes for each group. One-way ANOVA with Tukey’s multiple comparisons test was performed. (**g**) Signaling model for *Tlr2* induction. Recent reports indicate that the βγ subunit bound to Gi activates PLC in conjunction with Gq^58^. Inhibition of Gi by PTX inhibits this pathway. (**h**) qPCR analysis of effects of NA, carbachol, and inhibitors on expression of the *Relb* gene in isolated hepatocytes. The data are mean±s.d. *n*=3-5 biologically independent hepatocytes for each group. One-way ANOVA with Tukey’s multiple comparisons test was performed. (**i**) Signaling model for *Relb* induction.

To confirm direct action of NA on hepatocytes, we utilized isolated hepatocytes (Extended Data Fig. 9a and b). Hepatocytes expressed a broad repertoire of adrenergic and muscarinic receptors (Extended Data Fig. 9c). In isolated hepatocytes, NA—but not carbachol—triggered the induction of *Tlr2*, *Relb*, and *Saa1/2* (Extended Data Fig. 10a–c). These findings demonstrate that NA acts directly on hepatocytes, inducing the expression of *Tlr2*, *Relb*, and *Saa1/2* which are involved in the hepatocyte protective phenotype (Extended Data Fig. 10h).

To elucidate the regulatory mechanisms governing the NA-induced expression of *Tlr2* and *Relb* in hepatocytes, we used isolated hepatocytes. Adrenergic receptors (ARs) to which NA binds are coupled to heterotrimeric G proteins (Gs, Gi, and Gq/11), thereby linking to downstream PKA and PKC signaling pathways^33^. In isolated hepatocytes, NA-induced *Tlr2* expression was suppressed by inhibitors of PKA (H-89), Gi (pertussis toxin (PTX)), and PKC (Go6983) (Fig. 4f and g). Furthermore, the combined application of a PKA activator (dbcAMP) and a PKC activator (phorbol 12-myristate 13-acetate (PMA)) mimicked the effect of NA, inducing *Tlr2* expression (Fig. 4f and g). In contrast, *Relb* induction was independent of PKA but was significantly inhibited by PKC blockade (Fig. 4h and i). These results suggest that *Tlr2* expression requires the coordinated action of Gs-PKA and Gq-PKC pathways, whereas *Relb* induction is primarily driven by the Gq-PKC pathway (Fig. 4g and i). These are the first findings that NA regulates the expression of *Tlr2* and *Relb* in hepatocytes.

We next focused on the potential effects of NA on Kupffer cells. To investigate whether macrophage-specific genes were also regulated by NA to mediate inflammatory responses, we performed qPCR analysis on the liver samples from the wild-type mice. IL-6 and TNF-α promote hepatocyte proliferation, and ICAM-1 is involved in inducing the expression of these cytokines from Kupffer cells^34^. qPCR analysis showed that NA induced the expression of *Icam1*, *Il6*, and *Tnfa* in wild-type mice (Extended Data Fig. 7f, g, h). This induction was abolished following Kupffer cell depletion (clodronate). While *Icam1* and *Tnfa* were unresponsive to carbachol, *Il6* was induced by both NA and carbachol (Extended Data Fig. 7g, i, j, k, n). In the injury model, CCl_4_ also induced expression of *Icam1*, *Il6*, and *Tnfa*, which was suppressed by adrenergic antagonists (Extended Data Fig. 8g-l). Notably, in the 10-month-old *Mkk7* cKO mice, CCl_4_ failed to induce expression of *Icam1*, *Il6*, and *Tnfa* (Extended Data Fig. 8m-o). Together, these results suggest that the sympathetic nervous system (NA) promotes inflammatory responses by activating a transcriptional program in macrophages in the liver.

To confirm direct action of NA on macrophages, we utilized isolated macrophages (Extended Data Fig. 9a, b). Macrophages expressed a broad repertoire of adrenergic and muscarinic receptors (Extended Data Fig. 9c). In isolated macrophages, NA induced *Icam1* and *Il6* within 30 minutes (Extended Data Fig. 10d, e). While NA and carbachol showed additive effects on *Il6* induction, carbachol surprisingly suppressed the NA-induced expression of *Icam1* (Extended Data Fig. 10f, g). These findings demonstrate that NA acts directly on macrophages, inducing *Icam1* and *Il6*, which are involved in the inflammatory response. In contrast, carbachol induces *Il6* but antagonizes *Icam1* expression in macrophages (Extended Data Fig. 10h). Overall, these results show that NA and ACh regulate the expression of *Icam1* and *Il6* in macrophages, which promote hepatocyte proliferation.

The activation of hepatic stellate cells, which are also involved in liver regeneration and can be monitored by the marker *Acta2*, was induced by carbachol rather than NA (Extended Data Fig. 7l-n). *Acta2* expression during CCl_4_ injury was resistant to adrenergic blockade, suggesting regulation via the CCl_4_-ACh axis (Extended Data Fig. 8p, q). Thus, ACh regulates the expression of *Acta2* in hepatic stellate cells.

## Discussion

In this study, we show that the intrahepatic sympathetic network is rapidly activated upon sensing the damaged state of the liver. Based on these findings, we propose a novel “neuro-liver linkage model” of liver regeneration (Fig. 5). In this model, the CNS triggers a coordinated response via efferent sympathetic and parasympathetic pathways. Activated sympathetic nerves induce a transient surge of NA that governs hepatocyte metabolism and survival while priming Kupffer cells. Simultaneously, efferent parasympathetic nerves release a transient burst of ACh to activate Kupffer and hepatic stellate cells, thereby orchestrating the multicellular commencement of regeneration. In parallel, the adrenal glands—receiving sustained efferent sympathetic input from the CNS—produce a prolonged release of NA to regulate systemic circulation, including vasoconstriction and elevated blood pressure, to optimize physiological capacity at the organismal level. The nervous system thus provides an exquisite regulatory framework capable of coordinating diverse intrahepatic cell types while rapidly translating local signals into a synchronized systemic response.

**Figure 5.**
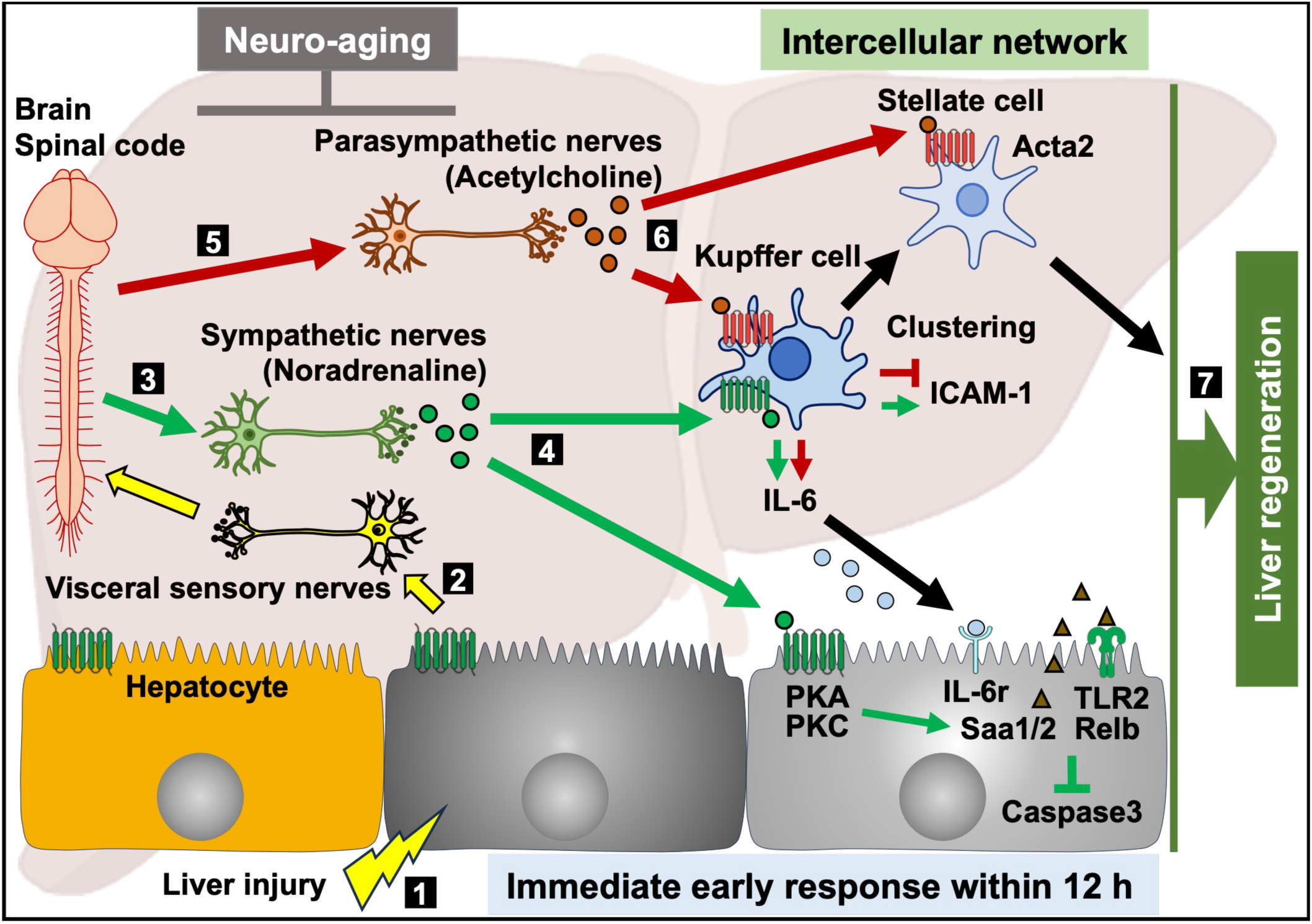
A proposed model of the autonomic nervous regulation of the initiation phase of liver regeneration. The scheme shows the immediate early response of the liver within 12 h after liver injury. 1. Liver injuries such as CCl_4_ and partial hepatectomy etc. 2. Injury signals released from the damaged hepatocytes stimulates the central nervous system of the spinal code and brain via visceral sensory nerves in the liver. 3. The sympathetic nervous system is activated. 4. Noradrenaline is released, transmitting signals to hepatocytes and Kupffer cells. In hepatocytes, noradrenaline activates PKA and PKC signaling to induce expression of genes, such as *Tlr2*, *Relb*, *Saa1/2*, and Caspase3 activation. In Kupffer cells, noradrenaline induces expression of genes, such as *Icam1* and *Il6* to form clustering and activate hepatocyte proliferation, respectively. 5. The parasympathetic nervous system is activated. 6. Acetylcholine is released, transmitting signals to Kupffer cells and hepatic stellate cells. These signals upregulate expression of genes, such as *Il6* in Kupffer cells and *Acta2* in hepatic stellate cells. In Kupffer cells, the acetylcholine attenuates *Icam1* expression. 7. Hepatocytes, Kupffer cells, and hepatic stellate cells cooperatively promote liver regeneration. Neuro-aging causes degeneration of both sympathetic and parasympathetic nerves, resulting in the impaired liver regeneration. Yellow arrows indicate injury signals. Green arrows are the sympathetic signals. Red ones are the parasympathetic signals. Black ones are the regeneration signals reported previously^2^.

The capacity of immune cells to synthesize and release ACh has gained attention. For instance, while splenic T-cell-derived ACh suppress macrophage activation^35^, recent evidence indicates that intrahepatic B cells produce ACh to conversely activate macrophages during liver regeneration^12^. Moreover, *in vitro* studies have shown that treating B cells with NA directly upregulates the expression of choline acetyltransferase (ChAT), the key enzyme for ACh synthesis^12^. However, the physiological source of this intrahepatic NA has remained elusive. Our study suggests that intrahepatic sympathetic-derived NA drives ACh production in B cells (Extended Data Fig. 11).

The age-associated attrition of the peripheral nervous system, characterized by axonal thinning and a progressive decline in innervation density, is a well-established hallmark of aging^36,37^. These degenerative changes are driven by a convergence of molecular failures, including dysfunctional motor proteins such as kinesin and dynein, cytoskeletal alterations involving microtubules and tau proteins, and compromised ATP production due to mitochondrial decay^38–43^. While a direct clinical link between MKK7 deficiency and peripheral nerve regression in humans has not yet been reported, our *Mkk7* cKO mouse model serves as a robust surrogate that faithfully recapitulates age-dependent axonal transport deficits and the resulting sarcopenia-like phenotypes observed in elderly populations. By leveraging this model, we have demonstrated that the dysfunction of the intrahepatic sympathetic–NA axis is a primary driver of the impaired regenerative capacity in the aging liver. Our findings suggest that age-related organ failure may be rooted not only in cell-intrinsic senescence but also in the systemic collapse of neural “information hubs” that coordinate rapid tissue repair. Contemporary longevity interventions have focused on targeting cell-intrinsic aging—such as utilizing senolytic drugs to clear senescent cells. However, restoring the nervous system could represent a promising therapeutic frontier for enhancing organ resilience and healthy aging. The concept of “inflamm-aging”, whereby chronic inflammation contributes to the aging process, has gained broad acceptance^44,45^. In addition, age-associated deterioration of neural function, termed “neuro-aging”, may constitute a fundamental mechanism underlying aging.

## Methods

### Animals

Mice carrying the *Mkk7*^flox^ or *Synapsin I-Cre* allele have been previously reported, respectively^46,47^. *Cre* recombinase expression under the *Synapsin I* promoter can be detected by embryonic day 12.5 in neuronal cells. Since the Cre recombinase driven by the *Synapsin I* promoter is expressed not only in the brain^5,31^, but also testis^48^, mice were prepared according to the crossing strategy as described in Extended Data Fig. 1b. The sperms of *Mkk7*^−/−^*Synapsin I-Cre* and *Mkk7*^flox/flox^ alleles were preserved in liquid nitrogen. Wild-type C57BL/6J male and female mice were obtained from Sankyo Labo or CREA Japan. All mice were housed in cages in a room on a standard 12 h light/ 12 h dark schedule and with access to standard food and water *ad libitum*. Mice were euthanized by inhalant anesthetic (isoflurane) overdose followed by cervical dislocation or decapitation. All procedures were conducted in accordance with ethical guidelines and regulations, and approved by the Institutional Animal Care and Use Committee of the Institute of Science Tokyo, and reported according to the relevant ARRIVE guidelines.

### Liver injury

Partial hepatectomy was performed as described previously^49^. In brief, mice were anesthetized using a mixture of medetomidine, midazolam, and butorphanol^50^. The abdomen was opened and the lateral and median lobes of the liver were ligated with sutures and resected. DDC treatment was performed by administering diet containing 0.1% DDC (CREA Japan) for 3 weeks. TAA treatment was performed by administering drinking water containing 300 mg/L TAA (33609-72, Nacalai Tesque) for 3 weeks. For the survival curve, 25% CCl_4_ (035-01273, FUJIFILM Wako Chemicals) in Olive oil (150-00276, FUJIFILM Wako Chemicals) was injected at 1 mL/kg body weight into the abdominal cavity twice a week for 8 weeks. Chronic liver injury by CCl_4_ was induced by administration of 25% CCl_4_ in Olive oil at 1 mL/kg body weight twice a week for 3 weeks. Acute liver injury by CCl_4_ was induced by single administration of 25% CCl_4_ in Olive oil at 1 mL/kg body weight.

### Hepatic vagotomy

Vagotomy was performed as described previously^9^. After anesthetized, the abdomen was shaved and disinfected with 70% EtOH. A longitudinal incision was made in the midline of the abdomen. The skin and peritoneum were detached with scissors. A longitudinal incision was made in the median peritoneum. The liver was turned over cephalad with a cotton swab. The connective tissue connecting the caudate lobe of the liver and the stomach was detached with tweezers. The esophagogastric junction was grasped with round-tipped tweezers and pulled caudally to facilitate viewing of the esophagus, and the anterior vagus nerve, which runs parallel to the esophagus, was identified. The hepatic branch of the vagus nerve branching from the anterior vagus near the lower esophagus was identified and cut sharply with scissors or bluntly with tweezers.

### Agonist and antagonist treatment

Carbachol (07136-14, Nacalai Tesque) and noradrenaline (A7256-1G, Sigma-Aldrich) were administered through the tail vein of mice at concentrations of 36.5 μg/mouse and 47.5 μg/mouse in 0.9% NaCl, respectively. Administration by drinking water containing 0.025 mg/mL prazosin (P0938, Tokyo Chemical Industry) or 0.0625 mg/mL propranolol (P0995, Tokyo Chemical Industry) started 24 h before CCl_4_ administration. Atropine (A0550, Tokyo Chemical Industry) was administered at 25 μg/g body weight in 0.9% NaCl solution intraperitoneally twice a day from one day before CCl_4_ treatment.

### Optical clearing of liver tissues

Optical clearing was performed as described previously^13^. Mouse livers were collected and fixed in PBS/1% paraformaldehyde (PFA)/10% sucrose for 1 h, followed by PBS/1% PFA for 2 h at room temperature. Tissues were washed three times in PBS for 1 h at room temperature, then washed overnight in PBS/30% sucrose. After washing three times in PBS for 1 h at room temperature, tissues were fixed in 20% methanol (diluted with distilled water) for 2 h, 40% methanol for 2 h, 60% methanol for 2 h, 80% methanol for 2 h, 100% methanol for 2 h, and incubated at room temperature. Tissues were decolorized with a mixture of 30% H_2_O_2_ and 100% methanol (v:v = 1:10) for 48 h at 4°C. Tissues were then shaken with 80% methanol for 2 h, 60% methanol for 2 h, 40% methanol for 2 h, and 20% methanol for 2 h, incubated overnight at room temperature in PBS/0.2% Triton X-100/0.1% deoxycholate. Tissues were blocked overnight with PBS/0.2% TritonX-100/5% normal goat serum (16210064, Gibco). Tissues were then immunolabeled with anti-tyrosine hydroxylase antibody (1:10, AB9702, Sigma-Aldrich) and anti-PGP9.5 antibody (1:10, 14730-1-AP, Proteintech) in PBS/0.1% Tween-20/5% normal goat serum for 72 h at room temperature. Tissues were washed in PBS/0.1% Tween-20 for 24 h at room temperature. Tissues were further immunolabeled with Alexa Fluor 488 goat anti-rabbit (diluted to a final concentration of 4 μg/ml, A11034, Invitrogen) in PBS/0.1% Tween-20/5% normal goat serum for 72 h. Tissues were washed in PBS/0.1% Tween-20 for 48 h. All incubations were performed with gentle rotation. Immunolabeled tissues were embedded in PBS/0.8% agarose blocks prior to the optical clarification step. Tissue blocks were incubated with 20% methanol (diluted with distilled water) at room temperature for 1 h × 2, 40% methanol for 2 h, 60% methanol for 2 h, 80% methanol for 2 h, 100% methanol for 1 h × 2, and 100% methanol overnight. Tissue blocks were incubated with a mixture of dichloromethane and methanol (v:v = 2:1) for 2 h, twice at room temperature, followed by three incubations with 100% dichloromethane for 1 h. Finally, the tissue blocks were incubated twice with 100% dibenzyl ether for 12 h at room temperature. All incubation steps were performed with gentle rotation. The length of neurons was measured using the Image J software (NIH) manually.

### Histological analysis

Dissected livers were fixed in 10% formaldehyde neutral buffer solution (3715251, Nacalai Tesque) overnight at 4°C with shaking, then incubated in 70% EtOH overnight. Then, they were paraffin-substituted using Thermo Excelsior ES. Paraffin blocks were prepared using paraffin-substituted livers. The prepared paraffin blocks were thin sliced at a thickness of 5 μm on a MICROM HM-325 (Thermo Fisher Scientific). The sections were deparaffinized with Xylene for 10 min × 2, 100% EtOH for 10 min × 2, 90% EtOH for 5 min × 1, and 70% EtOH for 5 min × 1, followed by a light wash in running water and a Milli-Q wash. For H&E staining, deparaffinized sections were stained with Mayer’s Hematoxylin (1.09249.0500, Sigma-Aldrich) for 30 sec to 1 min and treated with warm water at 42-45°C for 1 min. The sections were then stained with Eosin for 2 min. For immunostaining, deparaffinized sections were incubated in Target Retrieval Solution (pH6.1 S1699 or pH9.0, S2367 Dako) for 10 min at 110, blocked in 2.5% Normal Horse Serum (26050070, Gibco) at room temperature for 15 min, incubated with a primary antibody in TBS at 4 overnight, then incubated with ImmPRESS HRP reagent (MP-7401-50, Vector Laboratories) at room temperature for 30 min. After washing with TBS, sections were incubated with ImmPACT DAB solution (SK-4105, Vector Laboratories) for 3 to 5 min. The nucleus was stained with Mayer’s Hematoxylin. Antibodies used here were anti-Ki67 antibody (1:200, 652402, BioLegend), anti-F4/80 antibody (1:200, 70076, Cell Signaling Technology), anti-Cleaved Caspase-3 antibody (1:400, 9664, Cell Signaling Technology). Dehydration of the sections after staining was performed by the following procedure: 70% EtOH for 1 min × 1, 100% EtOH for 3 min × 2, 100% EtOH for 5 min × 1, and Xylene for 5 min × 2. Images were observed and photographed with a BZ-X710 microscope (KEYENCE). For the assessment of necrotic areas, liver tissues after H&E staining were used to quantify the pink-colored anucleated regions by Image J. For the assessment of hepatocyte proliferation and apoptotic cells, the number of Ki67- or Cleaved Caspase 3-positive cells was divided by that of hematoxylin-stained nuclei. For the assessment of macrophage numbers, F4/80-positive areas and whole tissue areas were quantified by Image J. Then, the ratio was calculated.

### Isolation of hepatocytes and macrophages

Isolation of primary mouse hepatocytes and liver non-parenchymal cells (LNPCs) was performed as described previously with slight modifications^51–53^. 8-11-week-old male mice were anaesthetized and subjected to *in situ* liver perfusion via the inferior vena cava with 25 mL of Liver Perfusion Medium (17701-038, Gibco) followed by 60 mL of Liver Digestion Medium (17703-034, Gibco), each at 37°C. The liver was excised, and the gallbladder was removed. The tissue was transferred to a 60-mm non-treated dish containing 4 mL warm Hepatocyte Wash Medium (17704-024, Gibco). After gently removing connective tissue and surface debris, the liver capsule was carefully peeled and hepatocytes and non-parenchymal cells (LNPCs) were released into the medium. The cell suspension was filtered through a 70 µm cell strainer (43-50070-01, pluriSelect) into a 50 mL conical tube pre-wetted with Hepatocyte Wash Medium. The original dish was rinsed with additional warm Hepatocyte Wash Medium (3 times), and the final volume was adjusted to 50 mL per liver. Cells were centrifuged at 20 × g for 3 min at room temperature (RT) with slow acceleration and deceleration. The supernatant (containing LNPCs) was collected, and the hepatocyte pellet was retained. For hepatocyte isolation, the pellet was resuspended in 10 mL Hepatocyte Plating Medium (DMEM low glucose (08456-36, Nacalai Tesque) supplemented with 5% FBS (171012, Nichirei Biosciences Inc) and penicillin-streptomycin (26253-84, Nacalai Tesque) per mouse. The suspension was mixed 1:1 with 90% Percoll (17089102, Cytiva) (9 mL Percoll + 1 mL 10 × PBS (11482-15, Nacalai Tesque) per mouse and centrifuged at 200 × g for 10 min at RT (slow acceleration/deceleration). The top layer was aspirated, leaving ∼1 mL above the pellet (live hepatocytes), which was resuspended in 20 mL of warm Hepatocyte Plating Medium. Cells were then centrifuged at 50 × g for 2 min at RT and passed through a 40 µm cell strainer (3-6649-01, AS ONE) into a pre-wetted 50 mL tube. Viable cells were counted using trypan blue exclusion and plated at 2 × 10 cells/well in collagen-coated 12-well plates (4815-010, IWAKI). Cells were cultured for 3 h at 37°C with 5% CO. After attachment, media were replaced with Hepatocyte Maintenance Medium (Williams E Medium (12551-032, Gibco) containing 2 mM L-glutamine (G8540, Sigma), and penicillin-streptomycin), and cells were stimulated with noradrenaline or carbachol for the indicated times. Cells were harvested in RLT buffer (74106, Qiagen) for downstream RNA extraction. For LNPCs isolation, cells were collected from the hepatocyte isolation supernatants. After the 20 × g spin, the supernatant was recentrifuged at 400 × g for 5 min at RT. The resulting pellet was resuspended in 5 mL of 27% Percoll and transferred to a 15 mL tube. A sterile Pasteur pipette was inserted to the bottom of the tube to carefully underlay 5 mL of 63% Percoll, generating a two-layer discontinuous gradient. Tubes were centrifuged at 2000 rpm for 20 min at RT with minimal acceleration and braking. The interphase containing enriched LNPCs was collected and washed with RPMI-1640 medium (30264-85, Nacalai Tesque) supplemented with 10% FBS and penicillin-streptomycin (RPMI+FBS). Cells were centrifuged at 1,500 rpm for 5 min, resuspended in 2–3 mL warm RPMI+FBS, and filtered through a 40 µm cell strainer into a fresh tube. The final volume was adjusted to 10–12 mL per mouse. Viable cells were counted and seeded at 7.5 × 10^5^ cells/well in 24-well plates (3820-024, IWAKI), followed by 2 h incubation at 37°C and 5% CO. After initial attachment, media were removed, and cells were washed three times with warm 1 × PBS (14249-24, Nacalai Tesque). Fresh RPMI+FBS was added and cells were cultured for 24 h. Cells were stimulated with or without compounds for the indicated times. Following stimulation, cells were harvested in RLT buffer for RNA extraction. H-89 (20 μM, HY-15979, MedChemExpress)^54^ and Go6983 (2 μM, HY-13689, MedChemExpress)^55^ were added 30 min before NA and carbachol treatment. dbcAMP (HY-B0764, MedChemExpress)^56^ and PMA (HY-18739, MedChemExpress)^55^ were used at 1 mM and 100 nM, respectively. For PTX treatment, mice were injected with PTX (168-22471, Fujifilm Wako) intraperitoneally at 200 ng/200 μL PBS 24 h before isolation^57^. After isolation, PTX was added to the culture medium at 100 ng/mL.

### RNA purification

Total RNA was purified from 100 mg livers or isolated cells by using 1 mL TRIzol^®^ Reagent (15596018, Invitrogen) or RNeasy Mini Kit (74106, Qiagen), respectively, according to the manufacture’s instruction. For RNA-seq analysis, purified RNA was treated with DNase and further purified by using RNeasy Mini Kit (Qiagen). RNA was electrophoresed on a 1.5% Agarose gel to identify 28S and 18S rRNA bands in a ratio of approximately 2:1.

### Quantitative real-time PCR

cDNA was synthesized from purified RNA by using ReverTra Ace^®^ qPCR RT Master Mix with gDNA Remover (FSQ-301, TOYOBO) according to the manufacture’s instruction. Quantitative real-time PCR reactions were performed by using THUNDERBIRD^®^ Next SYBR™ qPCR Mix (QPX-201, TOYOBO) on the CFX96 Real-Time System (Bio-Rad Laboratories, Inc.). PCR primer sequences used are listed in Supplementary Table 4.

### RNA-sequencing analysis

After total RNA purification, RNA-sequencing analysis was entrusted to TaKaRa Bio. In brief, SMART-Seq v4 Ultra Low Input RNA Kit for Sequencing (Clontech), SMART-Seq mRNA LP (TaKaRa Bio), Nextera XT DNA Library Prep Kit (Illumina), IDT for Illumina - DNA/RNA UD Indexes, Tagmentation (Illumina) were used to amplify double-stranded cDNAs and prepare the sequencing library. RNA-seq was performed using NovaSeq 6000 NovaSeq Control Software v1.7.5, Real Time Analysis (RTA) v3.4.4, and bcl2fastq2 v2.20. Sequence data were analyzed using DRAGEN Bio-IT Platform v3.7.5 or v4.3.6 (Illumina) with GRCm39 Release m28 as the reference sequence. Results are expressed in transcripts per million (TPM). Obtained data was analyzed on iDEP.96 (http://bioinformatics.sdstate.edu/idep96/).

### Measurement of neurotransmitters

For noradrenaline, 100 mg livers or adrenal glands were homogenated in 500 µL of PBS(-). Homogenates were centrifuged and the supernatant (300 µL) was used for the assay. Noradrenaline amounts were measured by using Norepinephrine ELISA Kit (KA1891, Abnova) according to the manufacture’s instruction. For adrenaline, 100 mg livers were homogenated in 500 µL of 0.01 N HCl. Homogenates were centrifuged and the supernatant (300 µL) was used for the assay. Adrenaline amounts were measured by using Epinephrine ELISA Kit (KA3837, Abnova) according to the manufacture’s instruction. For acetylcholine, 100 mg livers were homogenated in 500 µL of PBS(-). Homogenates were centrifuged and the supernatant (20 µL) was used for the assay. Acetylcholine amounts were measured by using EnzyChrom Acetylcholine Assay Kit (EACL-100, BioAssay Systems) according to the manufacture’s instruction.

### Statistical analysis

Quantification of images were performed with Image J or hybrid cell count software (BZ-H3C, Keyence, Japan). Analyses of statistical significance were performed using GraphPad Prism 7 or 8 (GraphPad Software Inc., San Diego, CA). Student’s *t*-test was used for comparisons between two groups. One-way analysis of variance (ANOVA) was used for comparisons among more than three groups. Two-way ANOVA was used for comparisons among more than three groups with two factors. Log-rank test was used for comparisons of survival rates. Data were considered statistically significant at *P* < 0.05.

## Acknowledgments

We would like to thank Prof. Shusaku Uchida (Institute of Science Tokyo), Emeritus Prof. Toshiaki Katada (The University of Tokyo), and the members of Nishina laboratory for their invaluable advice and Life Science Editors for editing assistance.

## Author Contributions

S.K. and H.Nishina conceived and design the study. H.Nagao, S.K., and K.D. performed experiments and acquired the data. H.Nagao, S.K., K.D., A.S., and H.Nishina analyzed the data. H.Nagao, S.K. and H.Nishina wrote the manuscript.

## Data Availability

The RNA-seq data have been deposited into Gene Expression Omnibus (GEO) under the number GSE329051 for Fig. 1f and GSE329278 for Fig. 4a will be available on Apr 1, 2027. All other data and materials used in this study are available from the corresponding authors (S.K. and H.Nishina) upon reasonable request.

## Declaration of generative AI technologies in the writing process

The author used ChatGPT, Gemini, and DeepL throughout the article as a tool to assist with grammar checks and refine expressions for clarity.

---

**Extended Data Figure 1. Characterization of *Mkk7* cKO mice.**

(**a**) Gene targeting of *Mkk7*^flox^ mice and detection of the recombinated region in the *Mkk7* gene by PCR. (**b**) Mating strategy to produce *Mkk7* cKO mice. Since the Synapsin I promoter works in the testis, male mice carrying the *Syn I-Cre* gene cannot be used for mating. (**c**) Confirmation of the *Mkk7* gene deletion by PCR. The *Mkk7* gene is deleted in the brain but not in the liver in *Mkk7* cKO mice. An unprocessed image was shown in Extended Data Fig.12a. (**d**) Western blot analysis of MKK7 proteins in the livers of *Mkk7*^flox/flox^ mice and *Mkk7* cKO mice at 10 months of age. The number indicates independent mice. Unprocessed images were shown in Extended Data Fig.12b. (**e**) Rota-rod analysis. *Mkk7* cKO mice at 10 months of age exhibited an age-related decline in motor performance due to motor neuron dysfunction. The data are mean±s.d. *n*=3 mice for each group. Two-way ANOVA with Sidak’s multiple comparisons test was performed.

**Extended Data Figure 2. Liver responses to chronic CCl_4_ treatment in *Mkk7* cKO mice.**

(**a**) Experimental scheme. (**b**) Measurement of liver injury markers in the serum at the indicated ages. The data are mean±s.d. *n*=3 mice for each group. Two-way ANOVA with Tukey’s multiple comparisons test was performed. (**c**) Heatmap of k-means-clustered differentially expressed genes in RNA-seq analysis of the livers of *Mkk7*^flox/flox^ mice and *Mkk7* cKO mice at 10 months of age after chronic CCl_4_ administration. *n*=3 mice for each group. Related to Fig.1f-i.

**Extended Data Figure 3. Intrahepatic sympathetic innervation of *Mkk7* cKO mice.**

(**a**) (Left) Staining of PGP9.5 in the left lobe of the liver from mice at 13 months of age. White arrows indicate neural fibers. (Right) Quantification of neural lengths. The data are mean±s.d. *n*=3 mice for each group. Unpaired two-tailed Student *t* test was performed. Related to Fig.2a, b and c. (**b**) Measurement of levels of NA and ACh in the livers of female wild-type mice at 8 weeks of age after CCl_4_ administration. The data are mean±s.d. *n*=4 mice for each group. One-way ANOVA with Tukey’s multiple comparisons test was performed. (**c**) Measurement of levels of NA and ACh in the livers of female *Mkk7*^flox/flox^ mice and *Mkk7* cKO mice at 13 months of age after CCl_4_ administration. The data are mean±s.d. *n*=2-4 mice for each group. Two-way ANOVA with Sidak’s multiple comparisons test was performed. (**d**) Scheme of *Mkk7* cKO mouse phenotypes.

**Extended Data Figure 4. Visceral sensory nerve signaling after liver injury.**

Signals of liver injury are sensed by visceral sensory nerves within the liver and transmitted to the central nervous system. Subsequently, signals are conveyed to the liver and adrenal glands via efferent sympathetic or parasympathetic nerves. Green and brown dots represent noradrenaline and acetylcholine, respectively.

**Extended Data Figure 5. Time-dependent histological changes of the liver until 24 h after CCl_4_ treatment.**

(**a**) Experimental scheme. (**b**) H&E staining, immunostaining with anti-F4/80 antibody and anti-Ki67 antibody of liver sections from wild-type mice at 10 weeks of age treated with CCl_4_ until 24 h. Black arrows indicate macrophage clusters. (**c**) Quantification of (b). The data are mean±s.d. *n*=3 mice for each group. One-way ANOVA with Tukey’s multiple comparisons test was performed.

**Extended Data Figure 6. Ontology analysis of genes whose expression was altered by noradrenaline and carbachol.**

(**a**) Unsupervised heatmap analysis of livers from wild-type mice at 9 or 10 weeks of age treated with NA or carbachol for the indicated time. (**b**) Ontology analysis of genes upregulated by NA compared to carbachol at 6 h after CCl_4_ treatment. Ontology analysis of genes upregulated by carbachol compared to NA at (**c**) 3 or (**d**) 6 h after CCl_4_ treatment. Related to Fig. 4a.

**Extended Data Figure 7. Quantitative PCR analysis of gene expression in the liver after treatment of noradrenaline and carbachol with or without clodronate.**

(**a**) Experimental scheme of treatment of clodronate and neurotransmitters. Expression levels of the *F4/80* gene. The data are mean±s.d. *n*=3 mice for each group. Two-way ANOVA with Sidak’s multiple comparisons test was performed. Expression of (**b**) the *Tlr2* gene, (**c**) the *Relb* gene by NA, and (**d**) the *Tlr2* gene, (**e**) the *Relb* gene by carbachol. Expression levels of (**f**) the *Icam1* gene, (**g**) the *Il6* gene, and (**h**) the *Tnfa* gene by NA and (**i**) the *Icam1* gene, (**j**) the *Il6* gene, and (**k**) the *Tnfa* gene by carbachol. Expression levels of the *Acta2* gene by (**l**) NA and (**m**) carbachol. The data are mean±s.d. *n*=3 mice for each group. Two-way ANOVA with Tukey’s multiple comparisons test (for (b)-(m)) was performed. (**n**) Summary of gene expression in response to neurotransmitters.

**Extended Data Figure 8. Quantitative PCR analysis of gene expression in the liver after CCl_4_ treatment in wild-type mice and *Mkk7* cKO mice.**

Expression levels of (**a**) the *Saa1/2* gene, (**c**) the *S100a8* gene, (**g**) the *Icam1* gene, (**i**) the *Il6* gene, (**k**) the *Tnfa* gene, and (**p**) the *Acta2* gene in the livers from wild-type mice at 8 weeks of age after CCl_4_ treatment. The data are mean±s.d. *n*=3 mice for each group. One-way ANOVA with Tukey’s multiple comparisons test was performed. Expression levels of (**b**) the *Saa1/2* gene, (**d**) the *S100a8* gene, (**h**) the *Icam1* gene, (**j**) the *Il6* gene, (**l**) the *Tnfa* gene, and (**q**) the *Acta2* gene in the livers of wild type at the 8 weeks of age in response to 12 h treatment of CCl_4_ and prazosin or propranolol. The data are mean±s.d. *n*=3 mice for each group. One-way ANOVA with Tukey’s multiple comparisons test was performed. Expression levels of (**e**) the *Saa1/2* gene, (**f**) the *S100a8* gene, (**m**) the *Icam1* gene, (**n**) the *Il6* gene, (**o**) the *Tnfa* gene in the livers of *Mkk7*^flox/flox^ and *Mkk7* cKO mice at 10 months of age in response to 12 h treatment of CCl_4_. The data are mean±s.d. *n*=3 mice for each group. Two-way ANOVA with Sidak’s multiple comparisons test was performed.

**Extended Data Figure 9. Isolation of hepatocytes and macrophages from the mouse livers.**

(**a**) Experimental scheme of isolation of cells. (**b**) Confirmation of isolated hepatocytes (Albumin) and macrophages (F4/80) by immunostaining. (**c**) The summary of expressions of adrenergic and cholinergic receptors in hepatocytes and macrophages examined by qPCR.

**Extended Data Figure 10. Gene expression analysis in the isolated hepatocytes and macrophages.**

Expression levels of (**a**) the *Tlr2* gene, (**b**) the *Relb* gene, and (**c**) the *Saa1/2* gene in isolated hepatocytes after treatment of NA and carbachol. The data are mean±s.d. *n*=4 biologically independent hepatocytes for each group. One-way ANOVA with Tukey’s multiple comparisons test was performed. Expression levels of (**d**) the *Icam1* gene and (**e**) the *Il6* gene in isolated macrophages after treatment of NA and carbachol. The data are mean±s.d. *n*=3-5 biologically independent macrophages for each group. One-way ANOVA with Tukey’s multiple comparisons test was performed. Expression levels of (**f**) the *Icam1* gene and (**g**) the *Il6* gene in macrophages after double treatment of NA and carbachol. The data are mean±s.d. *n*=4 biologically independent macrophages for each group. Two-way ANOVA with Sidak’s multiple comparisons test was performed. (**h**) Summary of effects of NA and acetylcholine on gene expression in isolated hepatocytes and macrophages (Kupffer cells).

**Extended Data Figure 11. A potential explanation of the NA source to activate intrahepatic B cells *in vivo*.**

Modares *et al*. previously demonstrated that following PHx, B-cell-derived ACh activates macrophages to stimulate IL-6 release, thereby promoting liver regeneration^12^. Although NA was postulated to be the key upstream factor driving this B-cell ACh production, its anatomical source has remained elusive. The injury-activated sympathetic nervous system uncovered in our study may serve as the primary origin of NA.

**Extended Data Figure 12. Unprocessed images.**

(**a**) An image of agarose gel electrophoresis related to Extended Data Fig.1c. (**b**) Western blots related to Extended Data Fig.1d.

## Extended Data Figure Methods

### Rotarod test

The Rota-Rod test was performed using a Mouse Rota-Rod (47650, Ugo Basile). Motor function was evaluated by measuring the time it took for the mouse to fall from a rotating rubber rod with a diameter of 3 cm. To ensure acclimation, mice were trained three times a day for four consecutive days, and the performance on the fourth day was evaluated. The rod was accelerated from 5 rpm to 20 rpm over the course of one minute. Subsequently, it was further accelerated to 45 rpm over the next minute and then maintained at 45 rpm for an additional minute.

### Measurement of hepatic injury markers

Blood was collected from mice anesthetized with isoflurane by orbital blood sampling. After standing at room temperature for 30 min, the blood was centrifuged at 3,000 × *g*. The supernatant was assayed for ALT and AST using SPOTCHEMTM EZ (SP-4430, arkray).

### Macrophage depletion

Clodronate Liposomes (F70101C-N, FormuMax) was administered intraperitoneally at 140□µg/100 µL/body 24 h before experiments.

### Western blot

The dissected liver was homogenated in the lysis buffer □50 mM Tris-HCl (pH7.5), 150 mM NaCl, 1 mM EDTA (pH8.0), 1% Triton X-100, 0.5% sodium deoxycholate, 0.1% SDS, 1 mM Na_3_VO_4_, 50 mM NaF, 1 mM phenylmethylsulfonyl fluoride, 5 μg/mL aprotinin, 20 μg/mL leupeptin, 20 μg/mL pepstatin A], incubated for 10 min at 4□ and centrifuged at 15 krpm at 4□. The supernatant was transferred to a new tube and its protein concentration was measured by using Pierce BCA Protein Assay kit. Proteins were then separated on a polyacrylamide gel by electrophoresis and transferred onto a PVDF membrane (Millipore). The membrane was blocked in 5% skim milk/TBST(0.1%) for 30 min at room temperature, and incubated with a primary antibody in Can Get Signal Solution 1 (TOYOBO) at room temperature for 1 h. The membrane was washed and incubated with a secondary antibody in Can Get Signal Solution 2 (TOYOBO) at room temperature for 1 h. The images were taken by Amersham Imager 680 system (GE Healthcare). Antibodies used here were anti-MKK7 antibody (1:1,000, ab52618, abcam) and anti-β-actin antibody (1:10,000, 66009-1-Ig, Proteintech), and the secondary antibodies were anti-Rabbit IgG, HRP-Linked F(ab’)_2_ Fragment from Donkey (1:6,000, NA9340-1ML, Cytiva), anti-Mouse IgG (H+L) from Goat (1:6,000, 115-035-146, Jackson ImmunoResearch).

### Immunocytochemistry

Cells were washed twice with PBS, fixed in 4% PFA in PBS for 15 min, and permeabilized with 0.2% Triton X-100 in PBS for 10 min. After blocking with 1% goat serum in PBS for 15 min, the cells were incubated with a primary antibody (diluted in 1% goat serum/PBS) at 4°C overnight. Following wash, the cells were incubated with an Alexa Fluor-conjugated secondary antibody in 1% goat serum/PBS, along with Hoechst 33342 (1:500, H3570, Invitrogen) to counterstain nuclei, at room temperature for 30 min. Antibodies used here were anti-Albumin antibody (1:200, A90-134A, Bethyl Laboratories) and anti-F4/80 antibody (1:200, 70076, Cell Signaling Technology), and the secondary antibodies were Alexa Fluor 555-conjugated anti-goat IgG (H+L) secondary antibody (1:300, A-21432, Invitrogen) and Alexa Fluor 647-conjugated anti-rabbit IgG (H+L), F(ab’)_2_ Fragment (1:300, 4414S, Cell Signaling Technology). Images were observed and photographed with a BZ-X710 microscope (KEYENCE).

