## Supplemental Figures for "The sympathetic nervous system initiates liver regeneration"

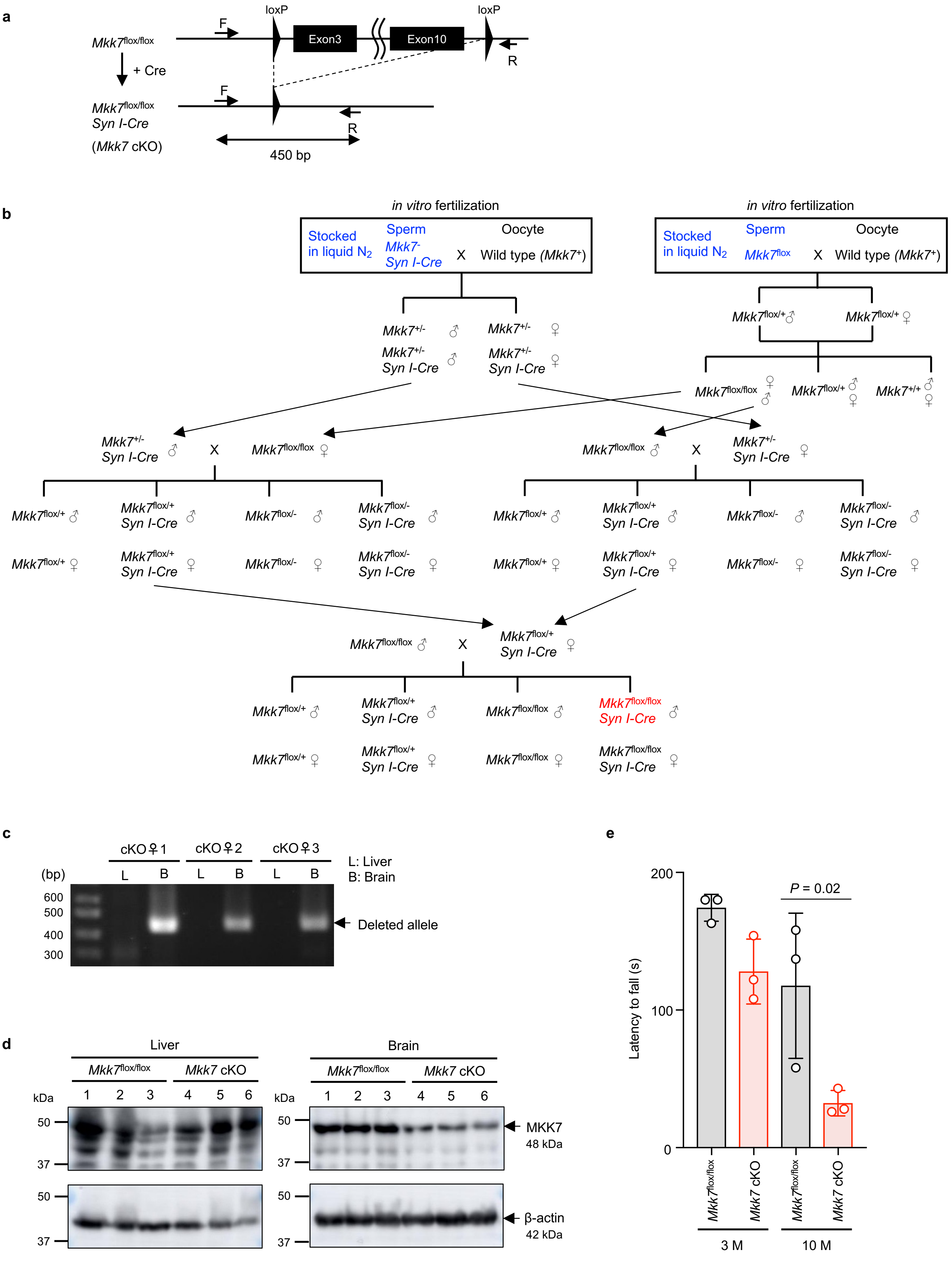


Extended Data Figure 1

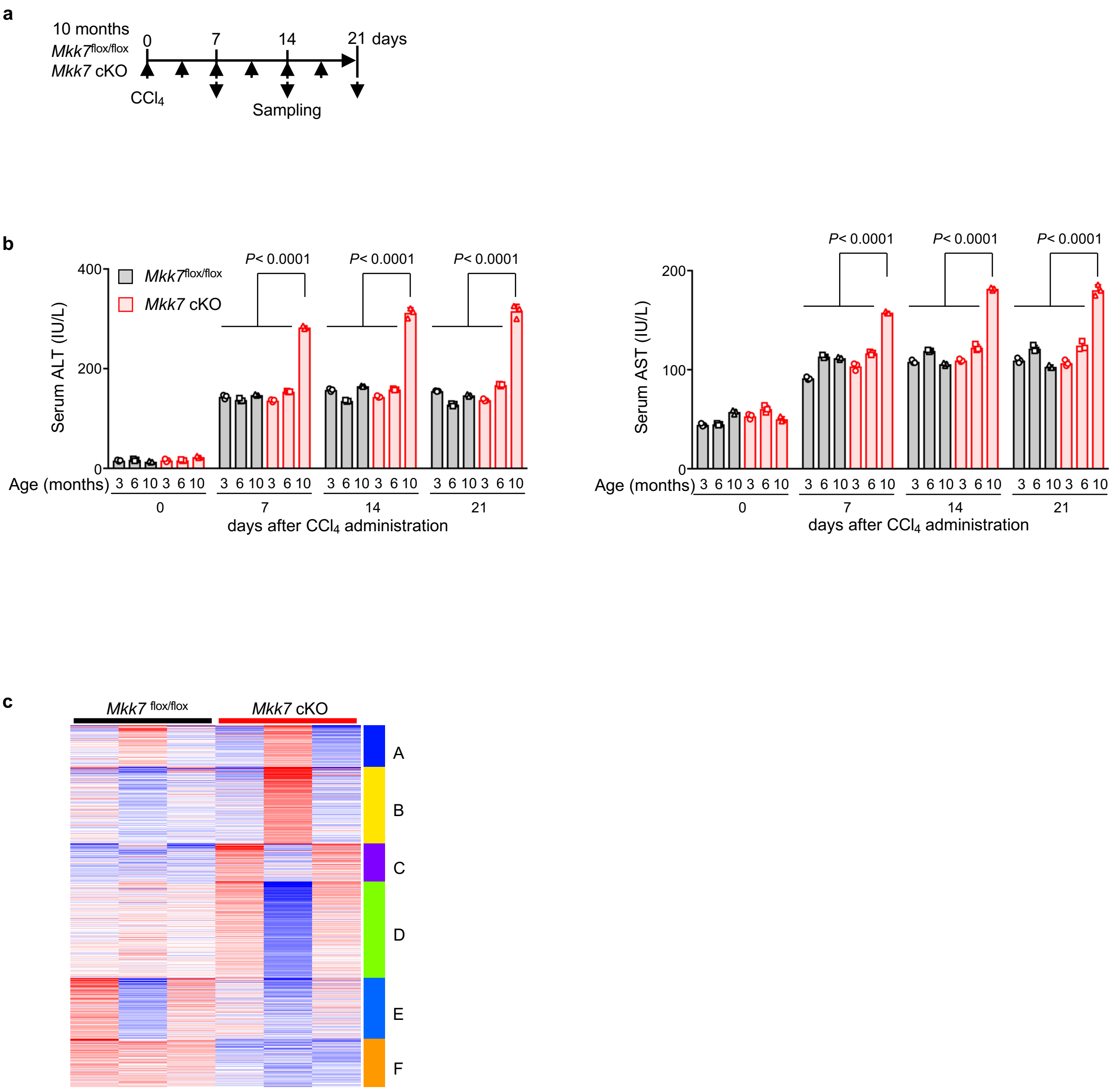


Extended Data Figure 2

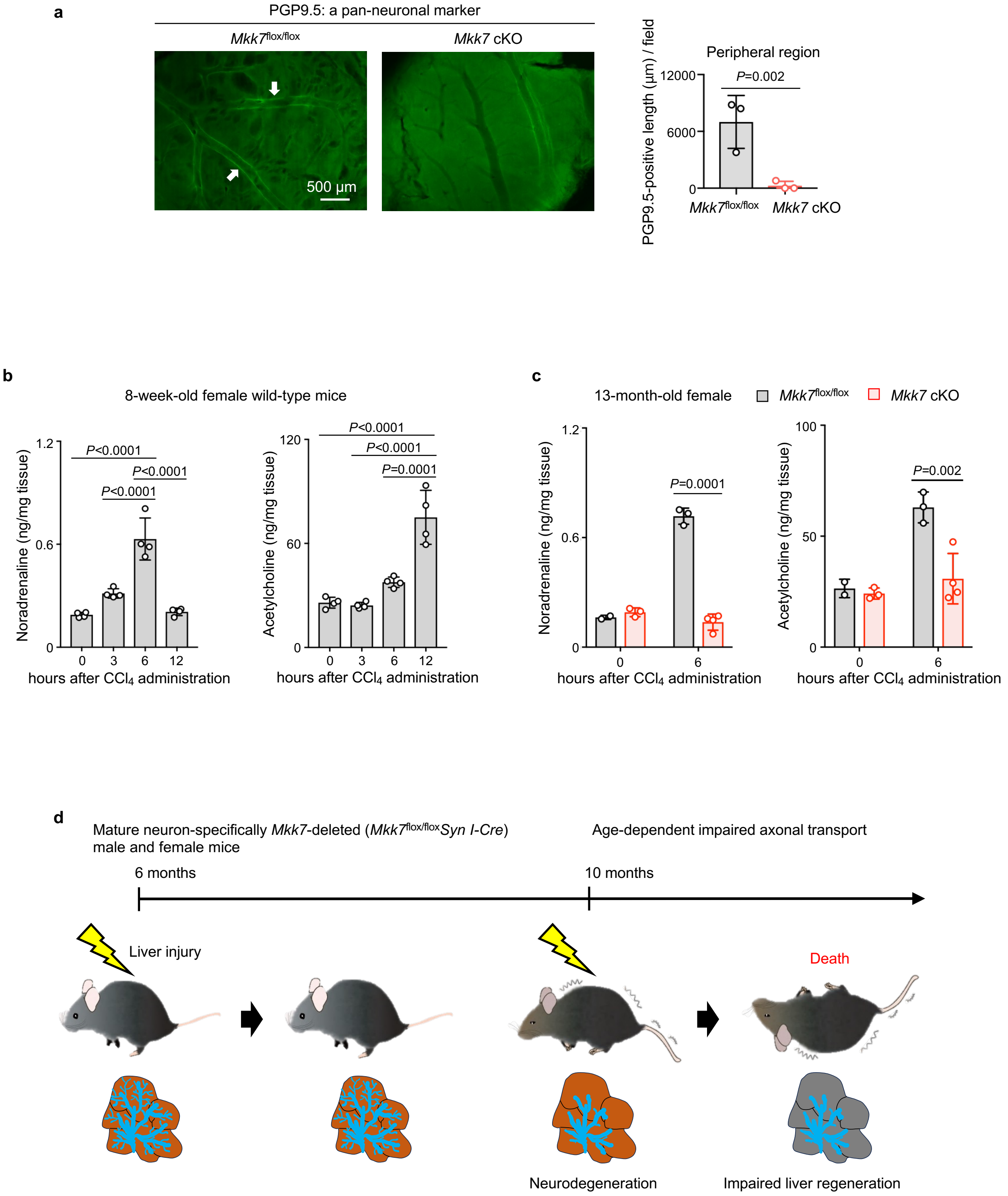


Extended Data Figure 3

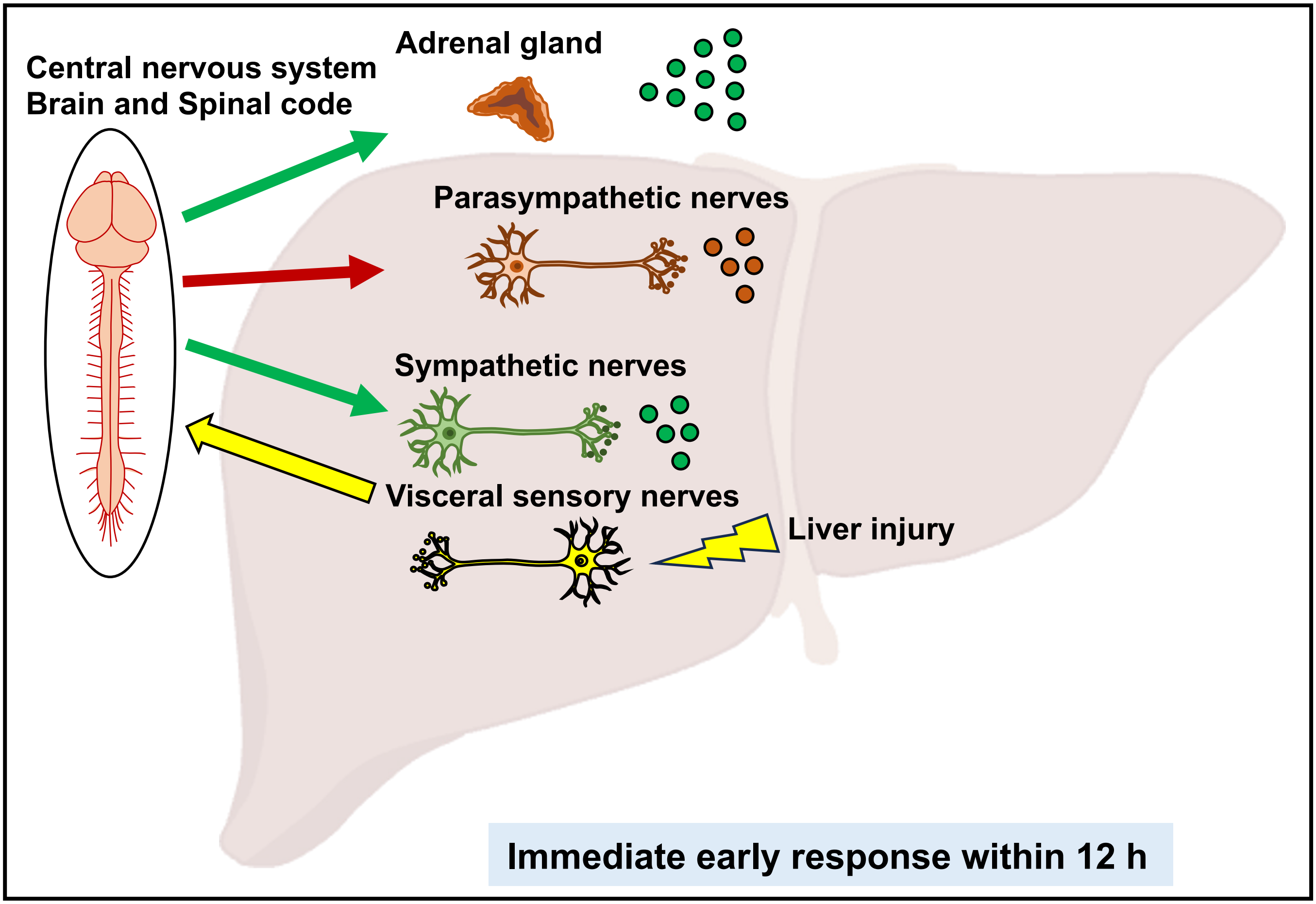


Extended Data Figure 4

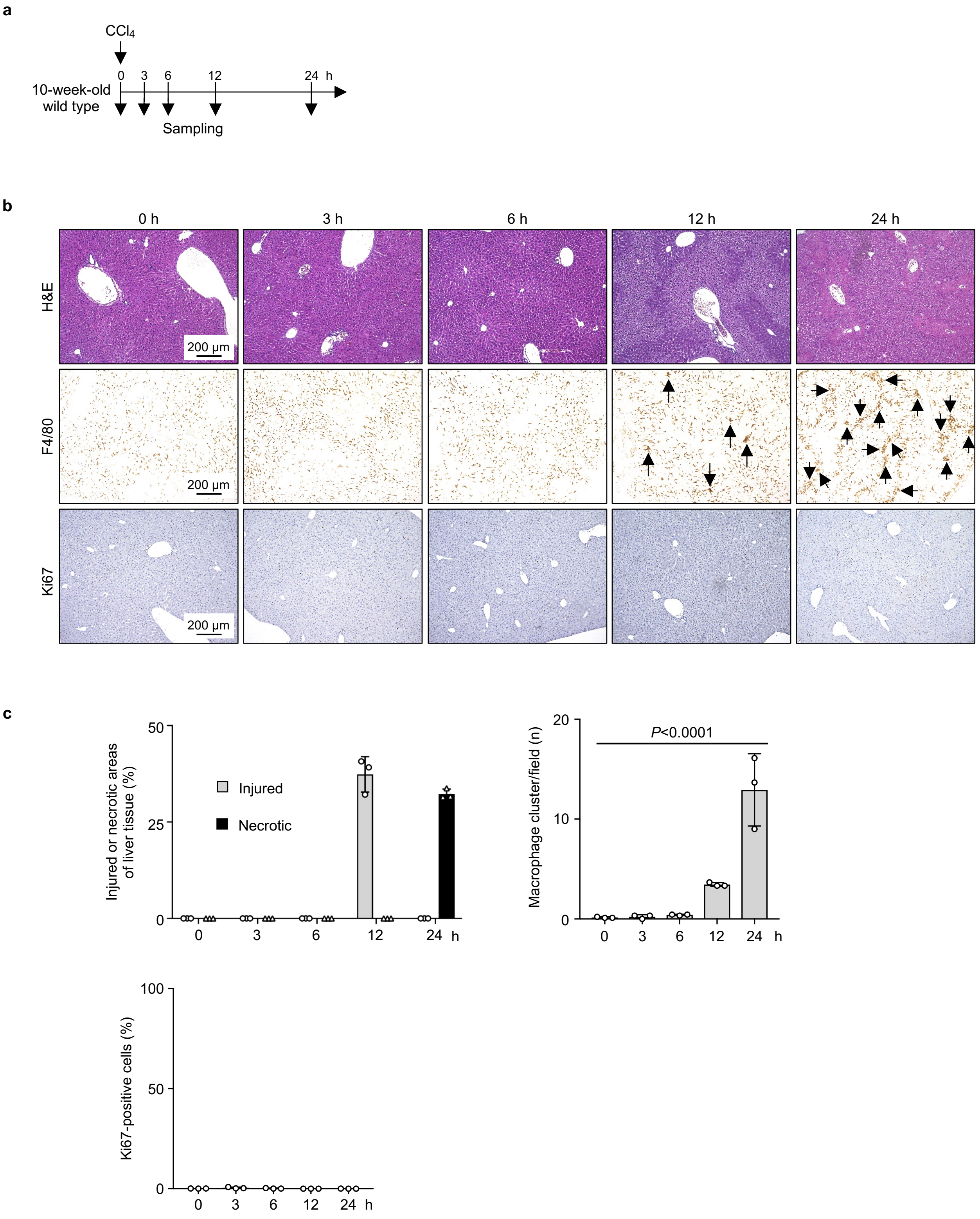


Extended Data Figure 5

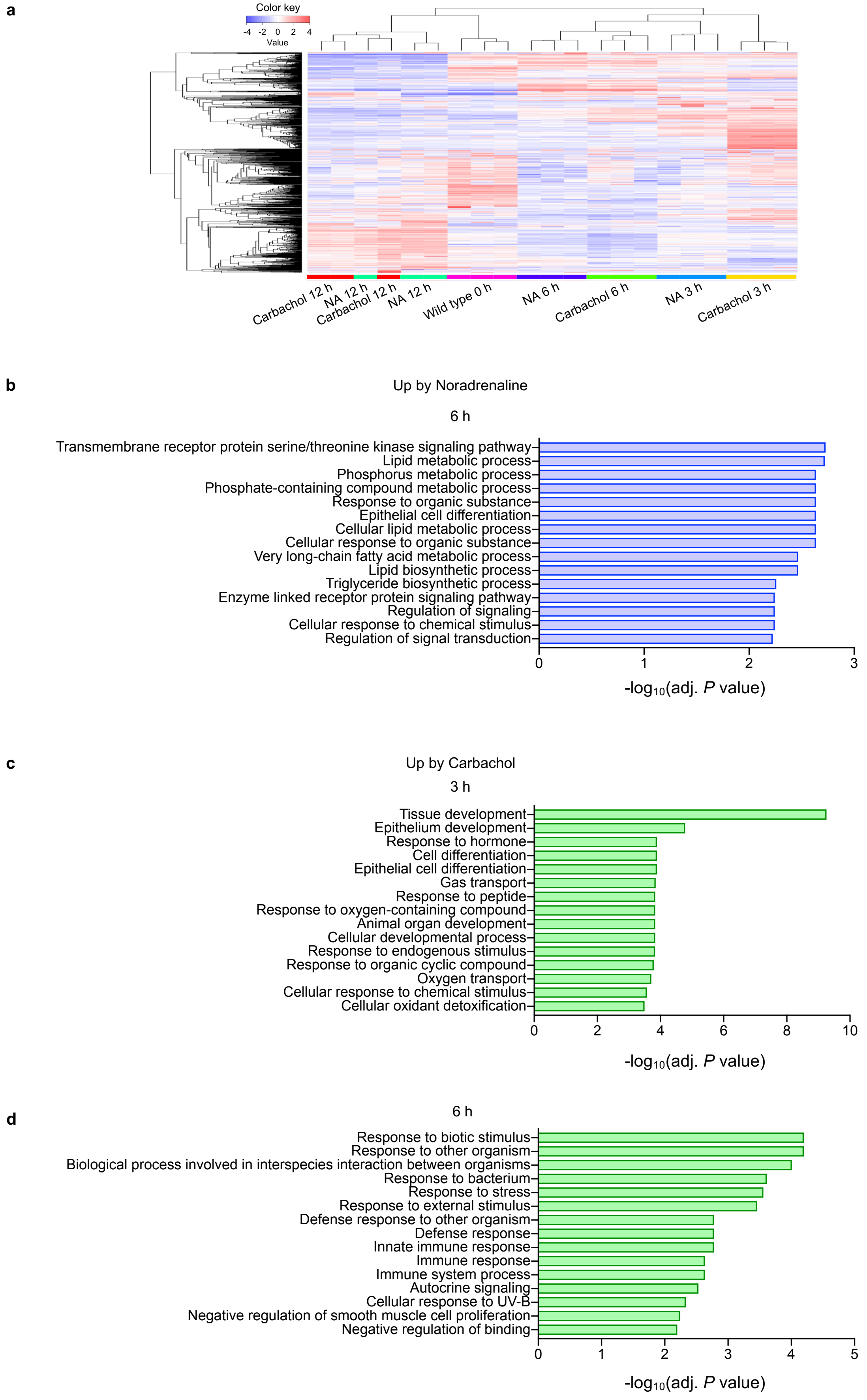


Extended Data Figure 6

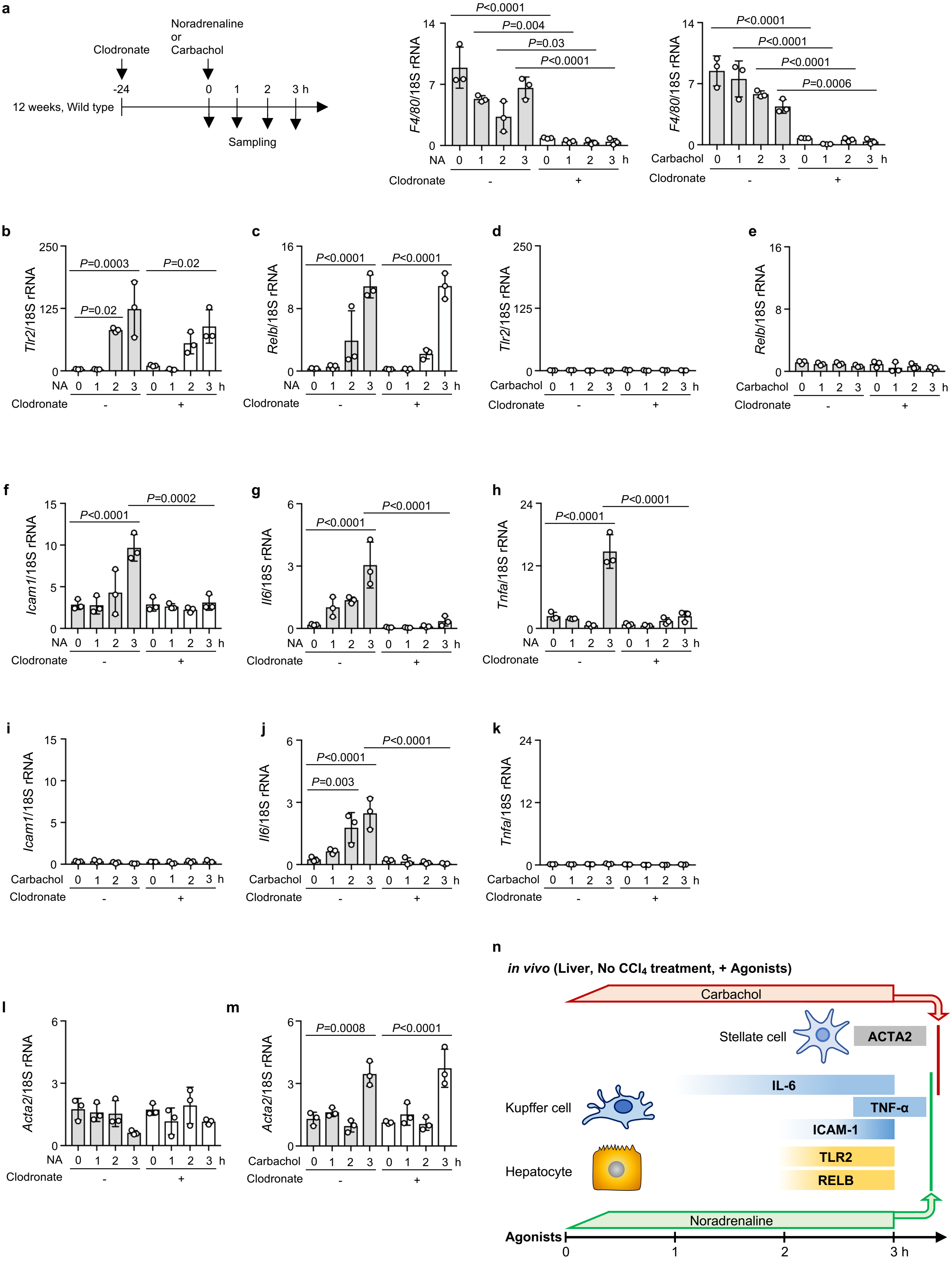


Extended Data Figure 7

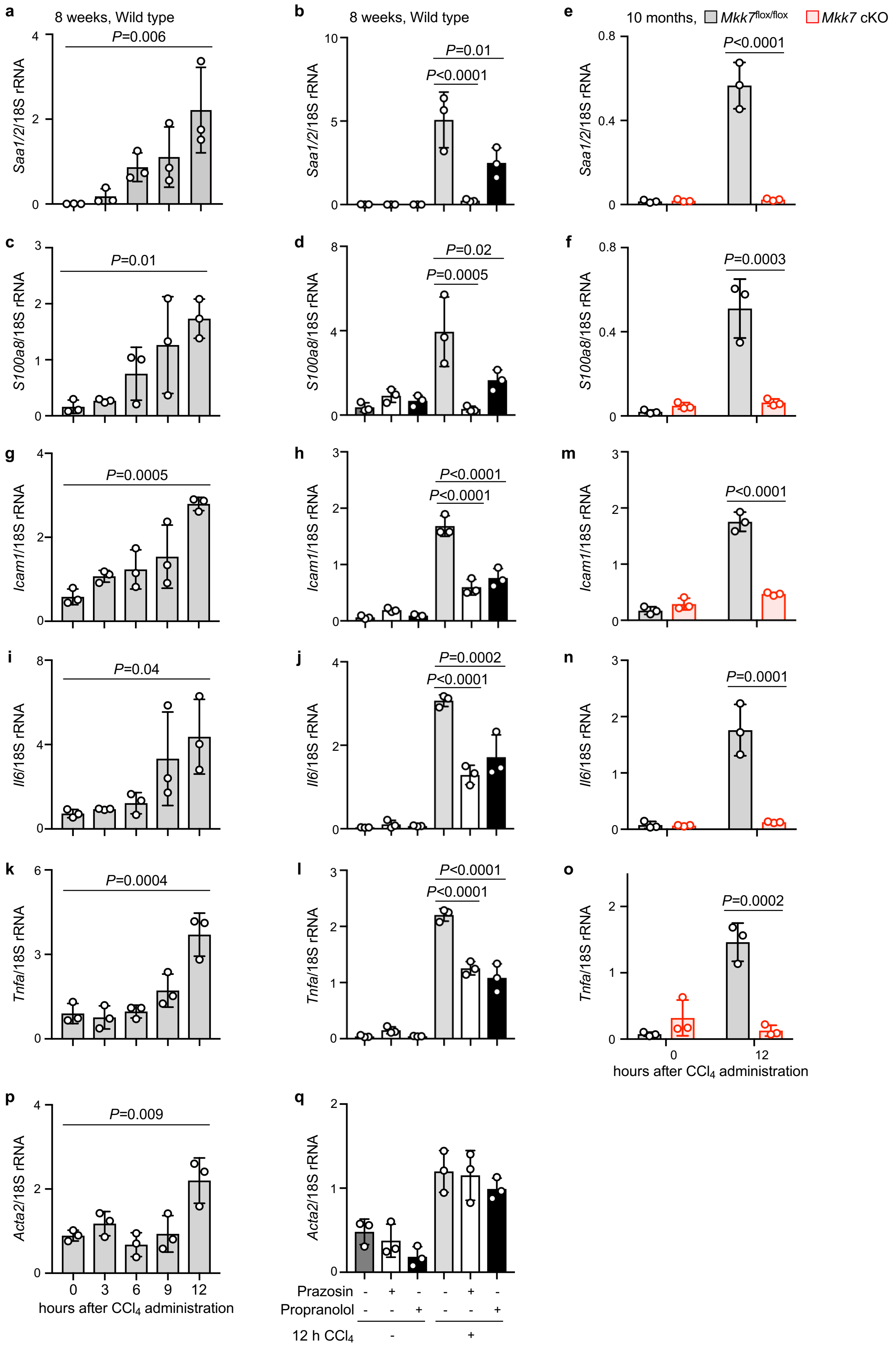


Extended Data Figure 8

a

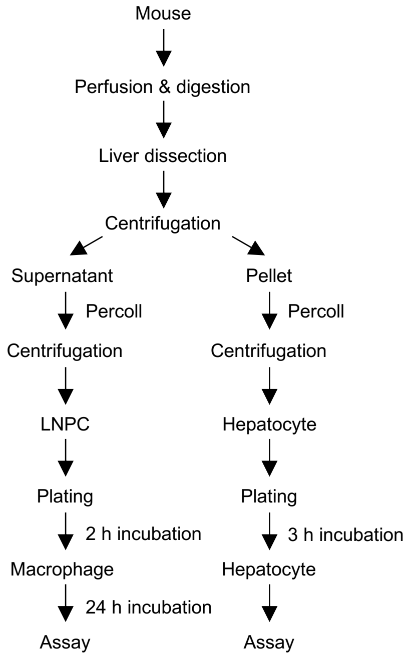


b

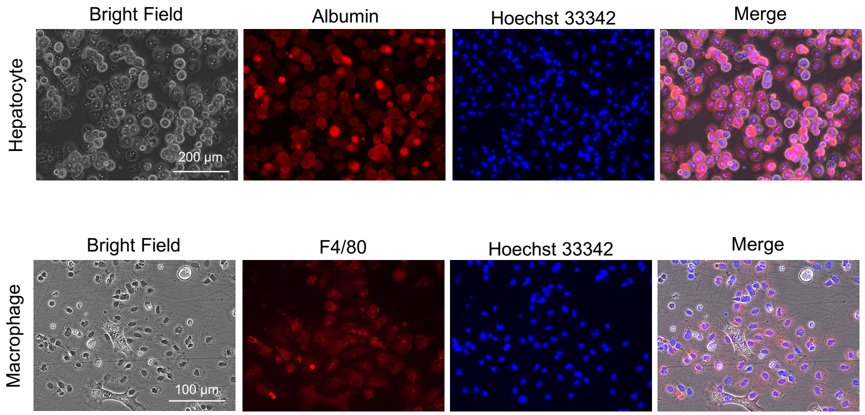


c

| | Receptor | G protein | Effector | Hepatocyte | Macrophage |
|---|---|---|---|---|---|
| Sympathetic | Adra1a | Gq/11 | PLC | + | + |
| | Adra1b | Gq/11 | PLC | + | + |
| | Adra2a | Gi/o | AC | + | + |
| | Adra2b | Gi/o | AC | + | + |
| | Adra2c | Gi/o | AC | +/- | + |
| | Adrb1 | Gs | AC | + | + |
| | Adrb2 | Gs | AC | + | + |
| | Adrb3 | Gs | AC | + | + |
| Parasympathetic | Chrm1 | Gq/11 | PLC | + | + |
| | Chrm2 | Gi/o | AC, K+ channel | +/- | + |
| | Chrm3 | Gq/11 | PLC | + | + |
| | Chrm4 | Gi/o | AC | + | + |
| | Chrm5 | Gq/11 | PLC | +/- | +/- |

Extended Data Figure 9

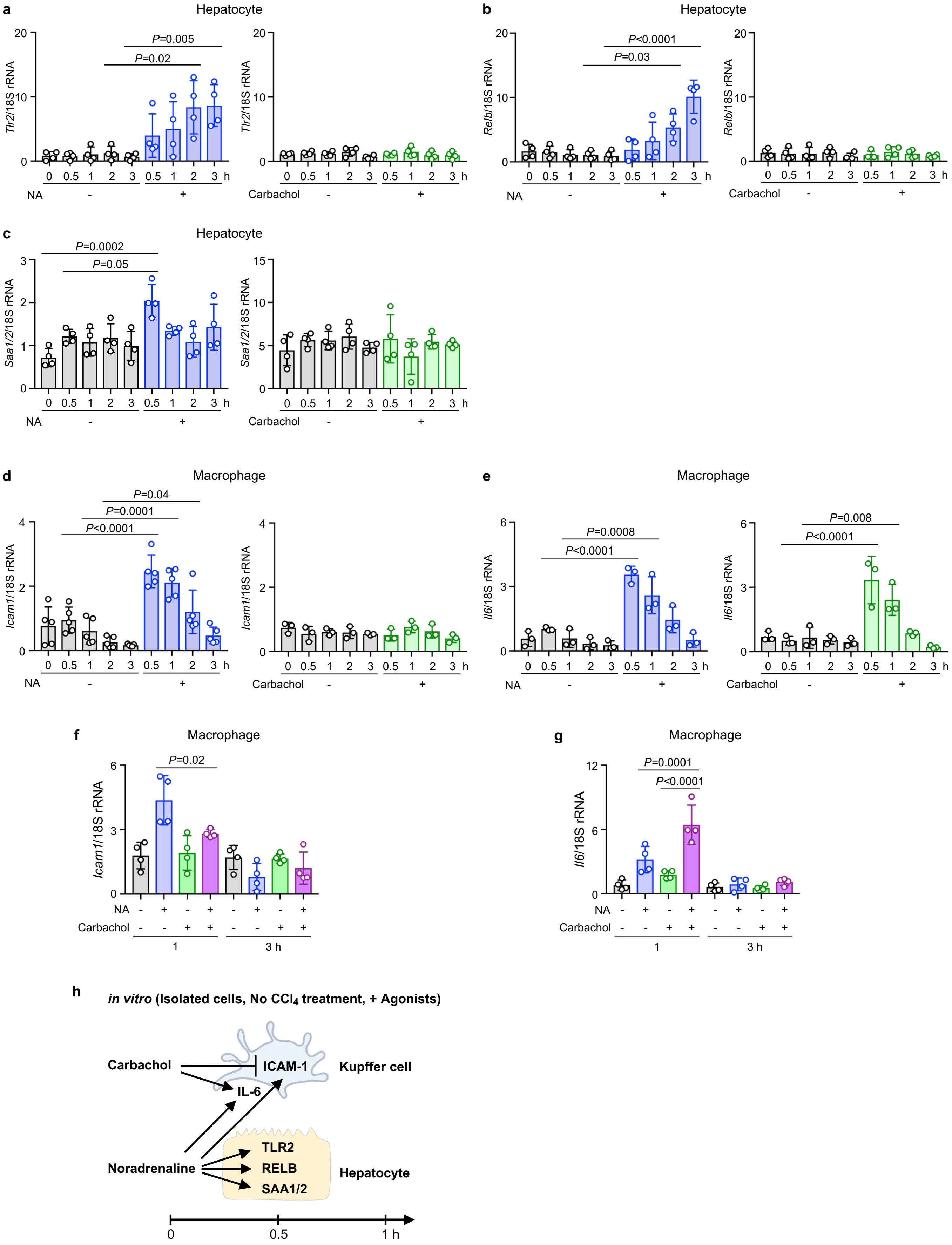


Extended Data Figure 10

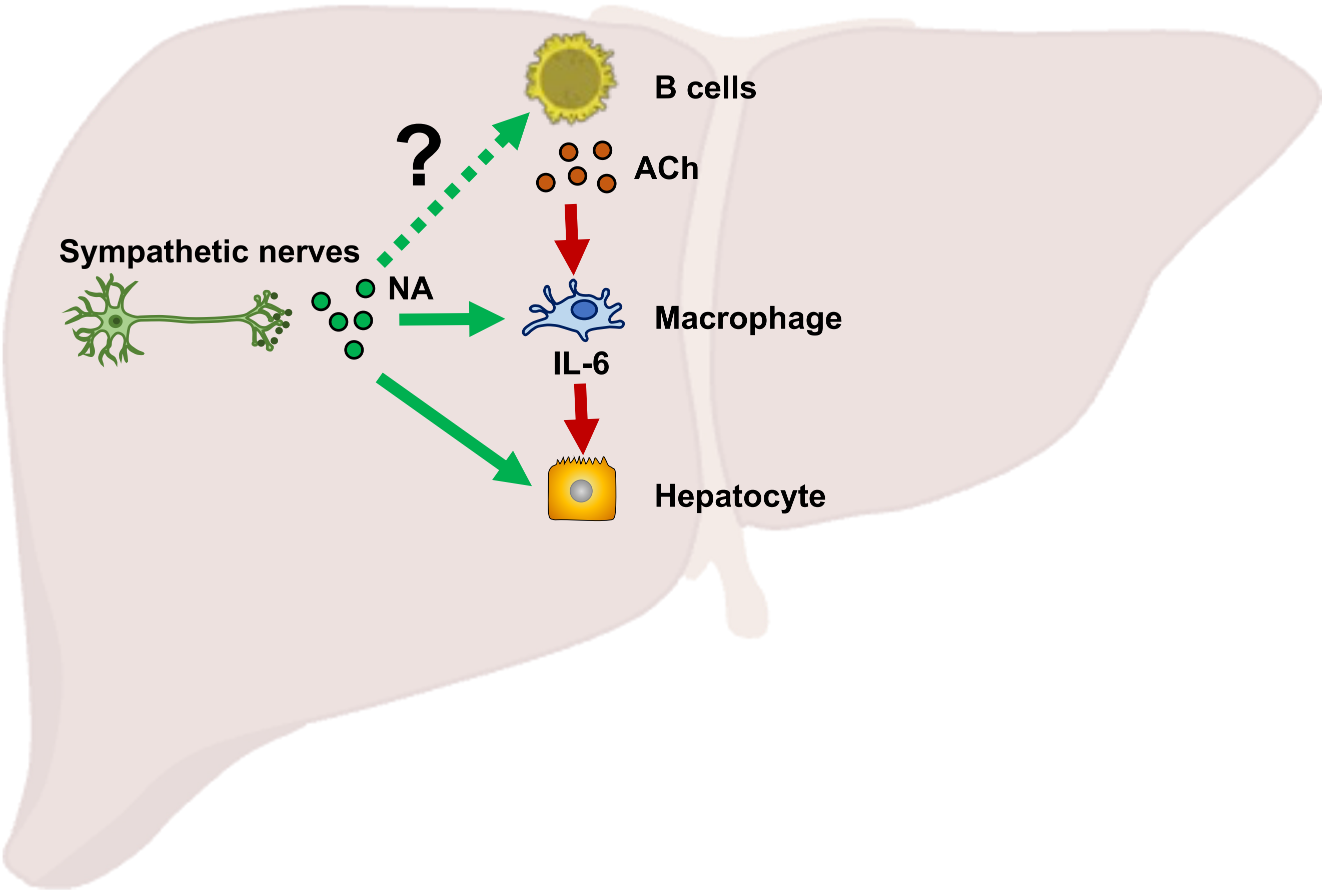


Extended Data Figure 11

a

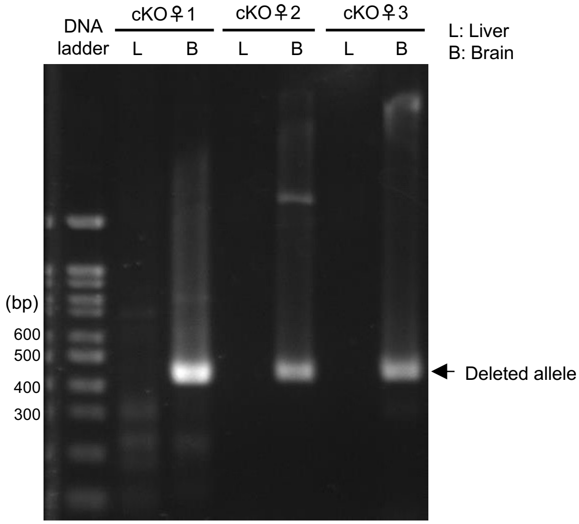


b

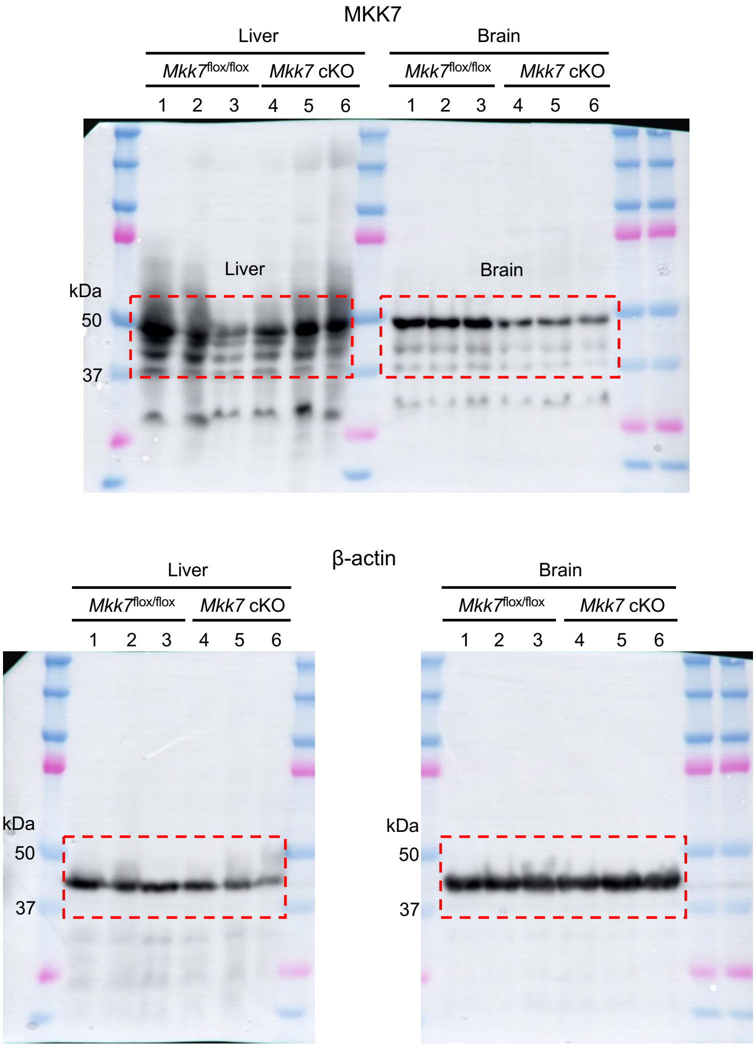


Extended Data Figure 12
